# Computational design of a protein family that adopts two well-defined and structurally divergent de novo folds

**DOI:** 10.1101/597161

**Authors:** Kathy Y. Wei, Danai Moschidi, Matthew J. Bick, Santrupti Nerli, Andrew C. McShan, Lauren P. Carter, Po-Ssu Huang, Daniel A. Fletcher, Nikolaos G. Sgourakis, Scott E. Boyken, David Baker

## Abstract

The plasticity of naturally occurring protein structures, which can change shape considerably in response to changes in environmental conditions, is critical to biological function. While computational methods have been used to de novo design proteins that fold to a single state with a deep free energy minima (Huang et al., 2016), and to reengineer natural proteins to alter their dynamics (Davey et al., 2017) or fold (Alexander et al., 2009), the de novo design of closely related sequences which adopt well-defined, but structurally divergent structures remains an outstanding challenge. Here, we design closely related sequences (over 94% identity) that can adopt two very different homotrimeric helical bundle conformations -- one short (∼66 Å height) and the other long (∼100 Å height) -- reminiscent of the conformational transition of viral fusion proteins (Ivanovic et al., 2013; Podbilewicz, 2014; Skehel and Wiley, 2000). Crystallographic and NMR spectroscopic characterization show that both the short and long state sequences fold as designed. We sought to design bistable sequences for which both states are accessible, and obtained a single designed protein sequence that populates either the short state or the long state depending on the measurement conditions. The design of sequences which are poised to adopt two very different conformations sets the stage for creating large scale conformational switches between structurally divergent forms.

## Introduction

There are many examples of de novo designed amino acid sequences that fold to a single designed target structure (Huang et al., 2016; Lu et al., 2018), but designing new amino acid sequences capable of adopting divergent structural conformations is challenging since the free energy difference between the two states must be small enough that a few amino acid changes can shift the global energy minimum from one state to the other, and both states must be stable relative to the unfolded state. Redesigns of natural protein backbones have yielded large conformational changes for systems such as the Zn antennafinger, Zinc-binding/Coiled coil (ZiCo), and designed peptide Sw2, that involve changes in oligomerization state from a homotrimeric 3-helix bundle to a monomeric zinc-finger fold (Ambroggio and Kuhlman, 2006; Cerasoli et al., 2005; Hori and Sugiura, 2004), and the pHios de novo design also involves a change in oligomerization state (pentamer to hexamer; (Lizatović et al., 2016)). In other cases, the engineered proteins have two well-defined states that are quite similar, for example the dynamic switching of DANCER proteins primarily involves a single tryptophan residue (Davey et al., 2017), and the Rocker channel has two symmetrically related states that are structurally identical (Joh et al., 2017). Finally, designed protein pH (Boyken et al., 2019) and peptide (Langan et al., 2019) dependent switches involve transitions between a single well-defined state and less well ordered states. Overall, in most previous work, the differences in conformation have generally either been relatively small, involved changes in oligomerization state, or only one of the two states has been well-defined. An exception is the engineering of sequences with 98% identity that switch folds between the naturally occuring G_A_ and G_B_ domains (Alexander et al., 2009).

We sought to design sets of closely related de novo protein sequences with two structurally divergent ground states having the same oligomerization state. Inspired by class I viral fusion proteins (i.e. influenza HA, parainfluenza F, HIV Env, and Ebola GP) (Ivanovic et al., 2013; Podbilewicz, 2014; Skehel and Wiley, 2000), we chose homotrimeric helical bundles as the fundamental architecture. We decided on a design scheme with two divergent conformations containing a constant six helix bundle “base”, and a variable portion with three inner helices and three “flipping” helices. In the compact “short” state (∼66 Å height) the flipping helices fold back to interact with the inner helices, and in the extended “long” state (∼100 Å height) they extend to form interactions with each other (Fig. 1a). Due to the difficulty of computing the relative stabilities of very different conformations accurately, we first designed related sequences which adopt one or the other of the two states. Then, guided by experiments, we identified structural features that affect the conformational preference between states and attempted to design single sequences which adopt both states.

**Fig. 1.**
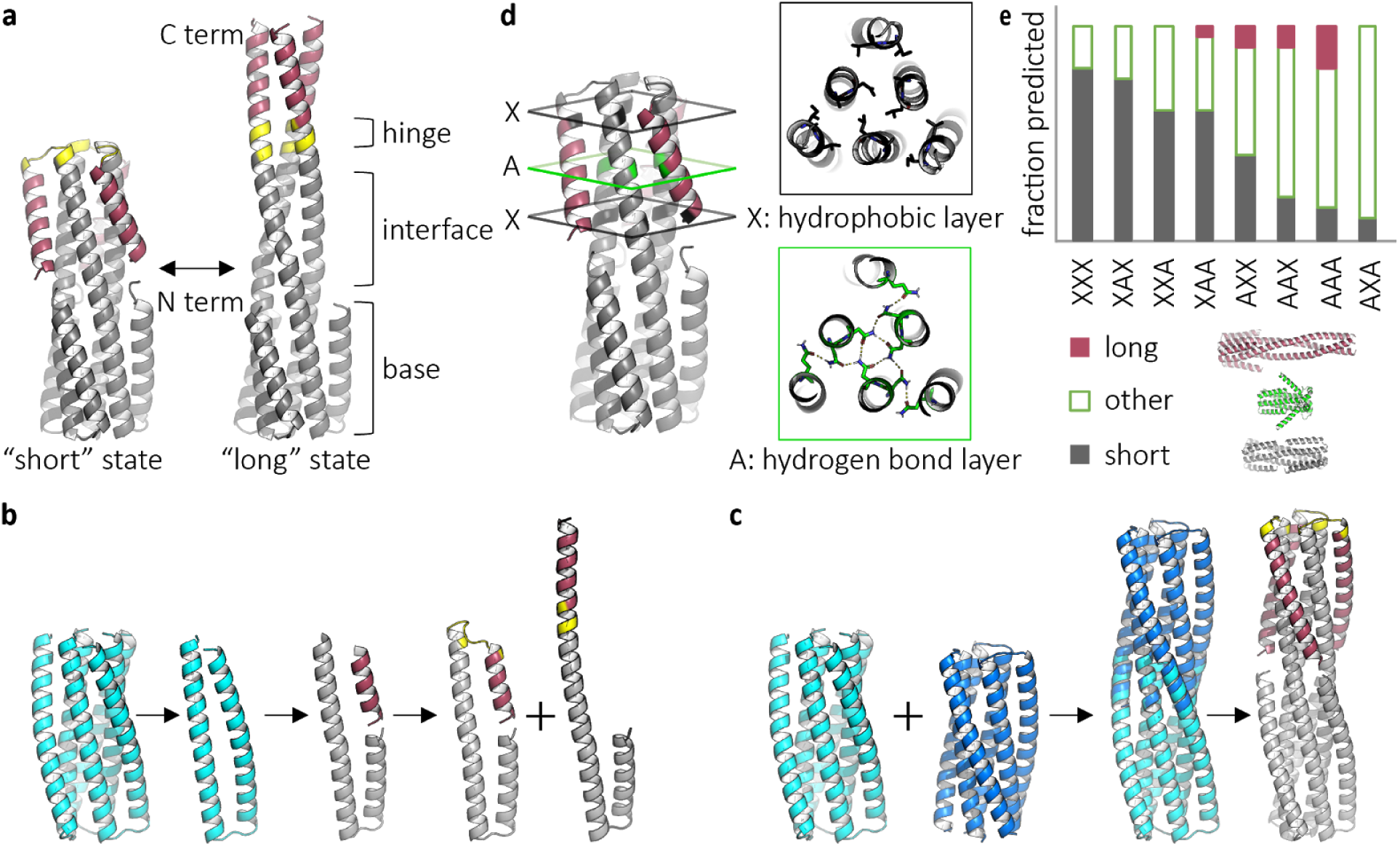
Design and in silico characterization of proteins with two distinct, well-defined ground states. **a**, Design concept for protein backbones with a “short” and “long” state. The backbone consists of a stable base, mobile flipping helices (red), tunable interface, and a flexible hinge (yellow). **b,** Initial design scheme. Starting with previously characterized protein 2L3HC3_13 (PDB ID 5j0h) variant (cyan), a monomer is extracted, cut to produce a flipping helix, and reconnected with a hinge to produce the backbone of the short or long state. **c,** Final design scheme. Previously characterized proteins 2L3HC3_13 variant (cyan) and 2L3HC3_23 (blue) are fused via their inner helices, outer helices are trimmed, and the hinge length is calculated such that the flipping helices pack against each other in the long state. **d,** The tunable interface between the inner helices and flipping helices. Each of three positions can be a hydrogen bond (green, “A”) or hydrophobic (black, “X”) layer. **e,** Fraction of top ten scored Rosetta folding predictions that resemble the short, long, or other structure for each permutation of possible interface configurations.

## Results

We started from a variant of a previously Rosetta-designed six-helix bundle (2L6HC3_13; PDB ID 5j0h) (Boyken et al., 2016) and introduced a cut in the outer helix to produce a disconnected segment to serve as the flipping helix. This flipping helix was connected to the C-terminus either via a turn to generate the “short” state, or a contiguous helix to generate the “long” state (Fig. 1b). To determine the appropriate hinge length, connectors of 2-7 residues were tested in silico and it was found that a hinge of length five provided good packing of the long state (Fig. S1b). Rosetta design and folding calculations were then used to find sequences that were strongly predicted to fold into one of the two states (Fig. S1d), genes encoding such designs were obtained, and the proteins were purified and characterized. All initial designs were solubly expressed, had circular dichroism (CD) spectra indicative of well-folded, alpha-helical structures, and three of four were trimeric by size exclusion chromatography with multi-angle light scattering (SEC-MALS) (Fig. S2). However, by small angle x-ray scattering (SAXS), it was difficult to determine which state (short, long, or a combination) the proteins adopted (Fig. S2).

To facilitate discrimination between the two states by SAXS, we carried out a second round of design with 1) longer flipping helices (which results in a greater difference in length between the states), and 2) helical interfaces redesigned to favor the short state (with sequence optimized for the turn and packing of the flipping helices on the inner helices) or the long state (with sequence optimized for packing between the flipping helices across the symmetric interface) — the previous design round enforced the same sequence at the interface regions for all designs (Fig. S3). The length of the flipping helix was increased from 15 to 21 residues and the optimal hinge length was recalculated to be three residues. The proteins in this second round of designs expressed solubly, and three of the four were helical by CD spectroscopy and ran as trimers by SEC-MALS (Fig. S4). However, by SAXS, all three appeared to be in the long state (Fig. S4), suggesting that the short state in this design scheme has higher free energy than the long state.

Our third round of designs aimed to generate short and long states with more equal stability. We used a larger portion of 2L6HC3_13 as the base to provide more stability, and fused it with a second Rosetta-designed six-helix bundle (2L6HC3_23) (Boyken et al., 2016), through aligning the matching superhelical parameters of the inner three helices (Fig. 1c). Compared to the second generation designs, in the short state, the flipping helix is positioned closer to the inner helices and the termini of the hinge region are closer together, allowing closer packing in the short state (Fig. S5). The outer helices were trimmed to avoid clashes and the required hinge length was recalculated to be three residues. Four designs with different fusion sites, placement of hydrogen bond networks, and loop flexibilities were tested; all expressed solubly and were helical by CD, but the two larger designs were not trimeric by SEC-MALS (Fig. S6). One design, AAA, was confirmed to be in the long state by crystallography, again indicating the short state has higher energy than the long state (Fig. 2f). We next focus on tuning the sequence of AAA to favor the short state.

**Fig. 2.**
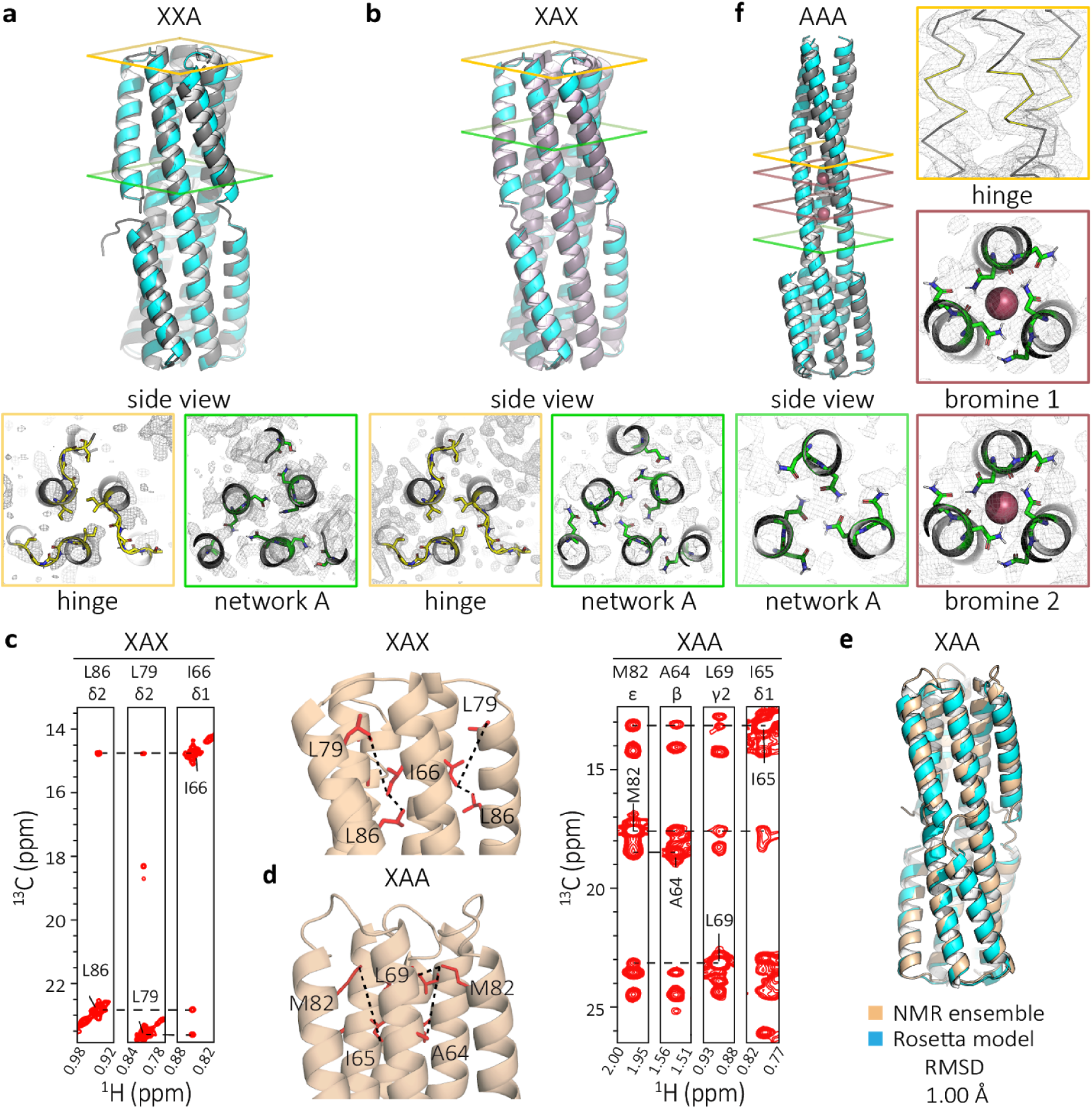
Modifying the number and position of hydrogen bond networks in the interface region changes the state. **a**, Crystal structure (grey) compared to model (cyan) of XXA (PDB ID 6nyi) shows the protein in the short state (RMSD = 1.37 Å). **b,** Crystal structure (grey) compared to the Rosetta model (cyan) of XAX (PDB ID 6nye) shows the protein in the short state (RMSD = 1.40 Å). **c,** 3D C_M_-C_M_H_M_ NOESY strip for XAX is consistent with the short conformation of the protein. **d,** 3D C_M_-C_M_H_M_ NOESY strip for XAA is consistent with the short conformational of the protein. **e,** De novo NMR structure (PDB ID 6o0i) calculated using chemical shifts, long-range NOEs and RDC data (beige) compared to the Rosetta model (cyan) shows the closed conformation (RMSD = 1.00 Å). **f,** Crystal structure (grey) compared to model (cyan) of AAA (PDB ID 6nx2,) show the protein in the long state with ion (red spheres) coordination sites (RMSD = 1.40 Å).

### Tuning the short/long state difference with hydrogen bond networks

To tune the relative accessibility of the short and long states, we used modular hydrogen bond networks that would be buried in the short state, but are partially exposed in the long state. The interface strength between the inner and flipping helices is expected to depend on both the number (more networks results in weaker interfaces) and position (solvent exposed locations are more likely to be disrupted by competing water interactions and result in weaker interfaces) of the networks. This strategy has the added advantage of creating more soluble proteins because residues necessarily exposed in the long state (when the flipping helix extends, parts of the inner helices it previously packed against are solvent exposed) are not entirely hydrophobic (as they would be if the stabilizing interactions were entirely non-polar, as in most design calculations). The interaction energy between the flipping helix and the inner helices in the short state was tuned by combining hydrogen bond (“A”) or hydrophobic layers (“X”) in different orders in each of three possible positions (Boyken et al., 2016) (Fig. 1d). Designs are given three letter names corresponding to the layers in order, starting at the hinge proximal position. Structure prediction calculations suggested that some layer combinations (such as XAX) are more favorable for the short state than others (such as AAA) and suggest that the short and long backbones states are among the lowest energy states of these systems (Fig. 1e).

We constructed the family of all eight possible permutations of two different layers in three positions, dubbed SERPNTs (in Silico Engineered Rearrangeable ProteiN Topologies). We set the three hinge residues to glycine to reduce favorable contributions to the long state free energy from the helical form of the hinge. All eight constructs expressed well, and by FPLC, most samples were monodisperse or had a small aggregation peak (only AXA formed a large soluble aggregate) (Fig. S7c). SEC-MALS measurements indicate that in solution, XAX, XAA, and AAA are trimeric as expected and CD wavelength and temperature measurements for XAX, XAA, and AAA show that the designs are helical and highly temperature stable (Fig. S7d,e).

The number and position of the hydrogen bond networks correlates with which state is lower energy -- designs with one network fold into the short state, two networks fold into a more dynamic short state, and three networks fold into the long state (details in supplementary text). Crystal structures of designs XXA (PDB ID 6nyi) and XAX (PDB ID 6nye) match the modeled short state (Fig. 2a,b). For both, most of the deviation, as expected, is in the flexible hinge: the crystal structures show that each monomer adopts a different loop. By NMR, the average solution structure adopted by XAX is locked in the compact state, as shown by the presence of multiple NOEs from a 3D C_M_-C_M_H_M_ NOESY spectrum (Fig. 2c). Analysis of an MAI(LV)^proS^ methyl-labeled XAA sample also suggests a short state, with the flipping helix shifted up (towards the hinge) compared to the design model (Fig. 2d,e, Fig. S8). The flipping helix appears to present a more flexible interface (relative to XAX), as evidenced by increased linewidths both in the amide TROSY and methyl HMQC spectra (Fig. S9 left). The crystal structure of design AAA (PDB ID 6nx2) is in the long state with some deviation from the predicted model because the asparagines of two of the hydrogen bond networks coordinate bromide ions and shift the helical register (ions and waters are not explicitly accounted for during modeling) (Fig. 2e, S10). These results suggested potential for a single sequence that can interconvert between the two designed states since a change in three residues, corresponding to the location of a hydrogen bond network, can switch the observed state (i.e., XAA vs AAA).

### Tuning the short/long state difference in the hinge region

To bring the amino acid sequence of the short and long states still closer together, we next tuned the hinge region, which transitions from a loop to fully helical in the conformational change (Fig. 2a,b,e vs f). In the long state, “key” hinge residue (G75) and the “backup” hinge residue (T76) have the potential to form an ion coordination site, but not in the short state (Fig. 3a). Similar natural sites include the calcium coordination site in human cytomegalovirus glycoprotein B (PDB ID 5cxf) or the nickel coordinating site in methylmalonyl CoA decarboxylase (PDB ID 1ef8) (Fig. 3b), among others (Fig. S11-S13) (Putignano et al., 2018). We sought to design calcium or nickel binding sites in the long state by introducing aspartic acid or histidine at the key hinge residue position. We also designed for structural ambiguity in the turn -- with the result that secondary structure prediction for residue 75 is sometimes helical and sometimes coil, but with low confidence (Fig. S14).

**Fig. 3.**
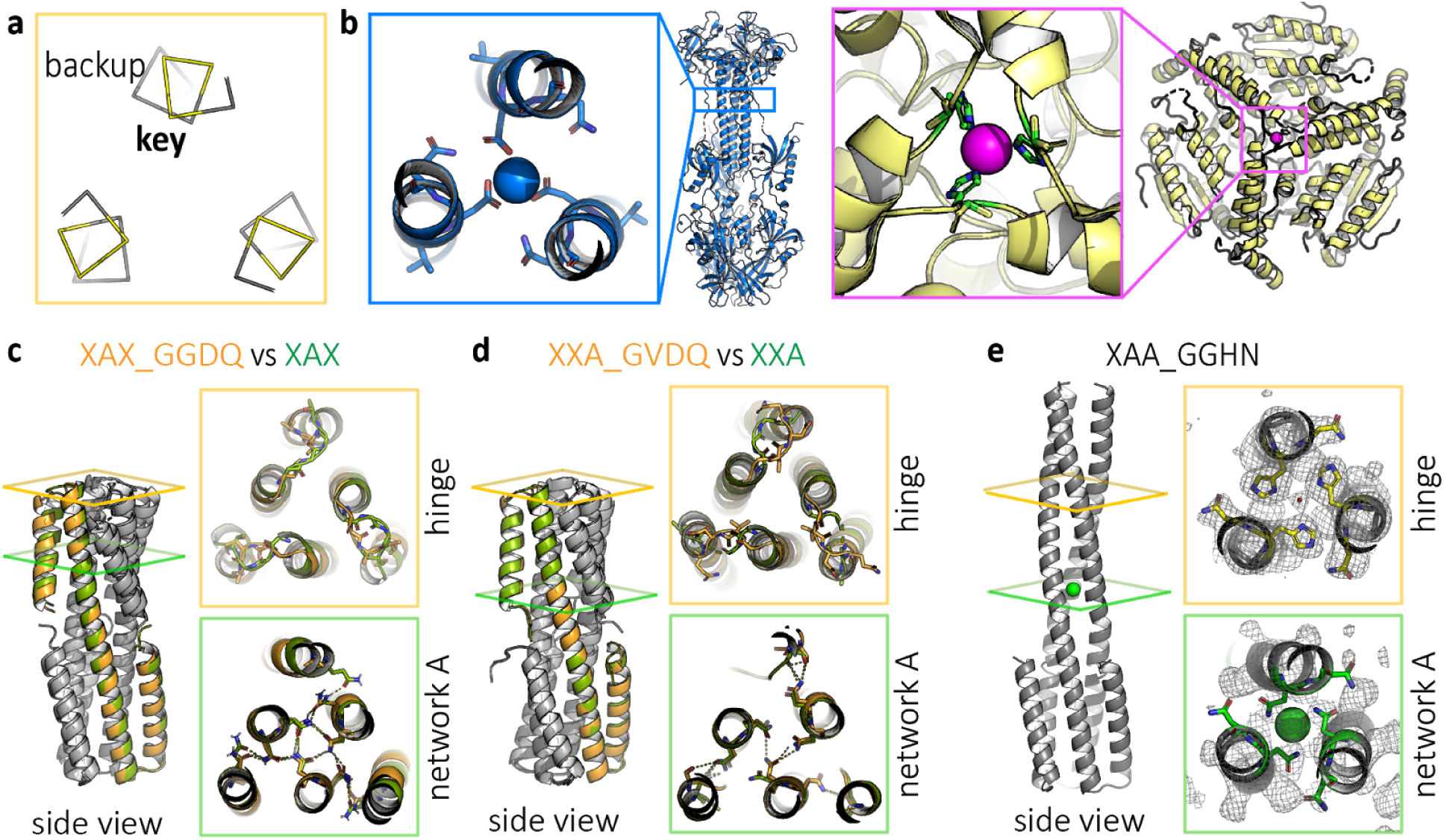
Tuning the hinge region can change the protein state. **a**, Position of key hinge residue G75 and backup hinge residue T76 have the potential to form an ion coordination site in the long state, shown here, but not in the short state. **b,** Examples of natural sites include calcium coordinating residues of human cytomegalovirus glycoprotein B (PDB ID 5cxf, blue) and nickel coordinating residues of methylmalonyl CoA decarboxylase (PDB ID 1ef8, yellow). **c,** Crystal structure of hinge modification XAX_GGDQ (orange, PDB ID 6nyk) is in the short state, like it’s parent XAX (green). **d,** Crystal structure of hinge modification XXA_GVDQ (orange, PDB ID 6nz1) is in the short state, like the parent XXA (green). **e**, Crystal structure of hinge modification XAA_GGHN (PDB ID 6nz3) is in the long state, unlike parent XAA.

Crystal structures show that designs XAX_GGDQ (PDB ID 6nyk) and XXA_GVDQ (PDB ID 6nz1), which introduces a potential calcium coordination site, adopt the short state like their respective base designs without the hinge residue modifications (i.e., XAX and XXA) (Fig. 3c,d). In contrast, the crystal structure of XAA_GGHN (PDB ID 6nz3), which introduces a potential nickel coordination site (occupied by a water molecule in the structure), is in the extended state, unlike the parent XAA design, with RMSD 1.44 to the extended model (Fig. 3e). Together, these results show that the hinge region is amenable to mutation without causing the protein to misfold into undesigned conformations.

### A bistable designed sequence that adopts both the short and long states

Having found that both the short and long state backbones are compatible with modular replacement of hydrogen bonding layers between the helices and with mutations in the hinge region, we carried out more detailed structural characterization to determine if any of the designs can adopt both conformations. Detailed NMR and crystallographic characterization of design XAA_GVDQ, described in the following paragraphs, suggest that this sequence may indeed populate both the short and long states.

To characterize the structure in solution, we prepared a selective MAI(LV)proS methyl-labeled XAA_GVDQ sample (Fig. S9 right). In the absence of calcium, the 3D CM-CMHM NOESY and residual dipolar couplings (RDCs), are consistent with the short state. We solved the NMR structure using the chemical shifts, long-range NOEs (both methyl to methyl and amide to methyl), and RDCs, and found it to be very close to the short state design model, RMSD 1.19 Å (Fig. 4a,c, Fig. S15, Table S6, PDB ID 6O0C).

**Fig. 4.**
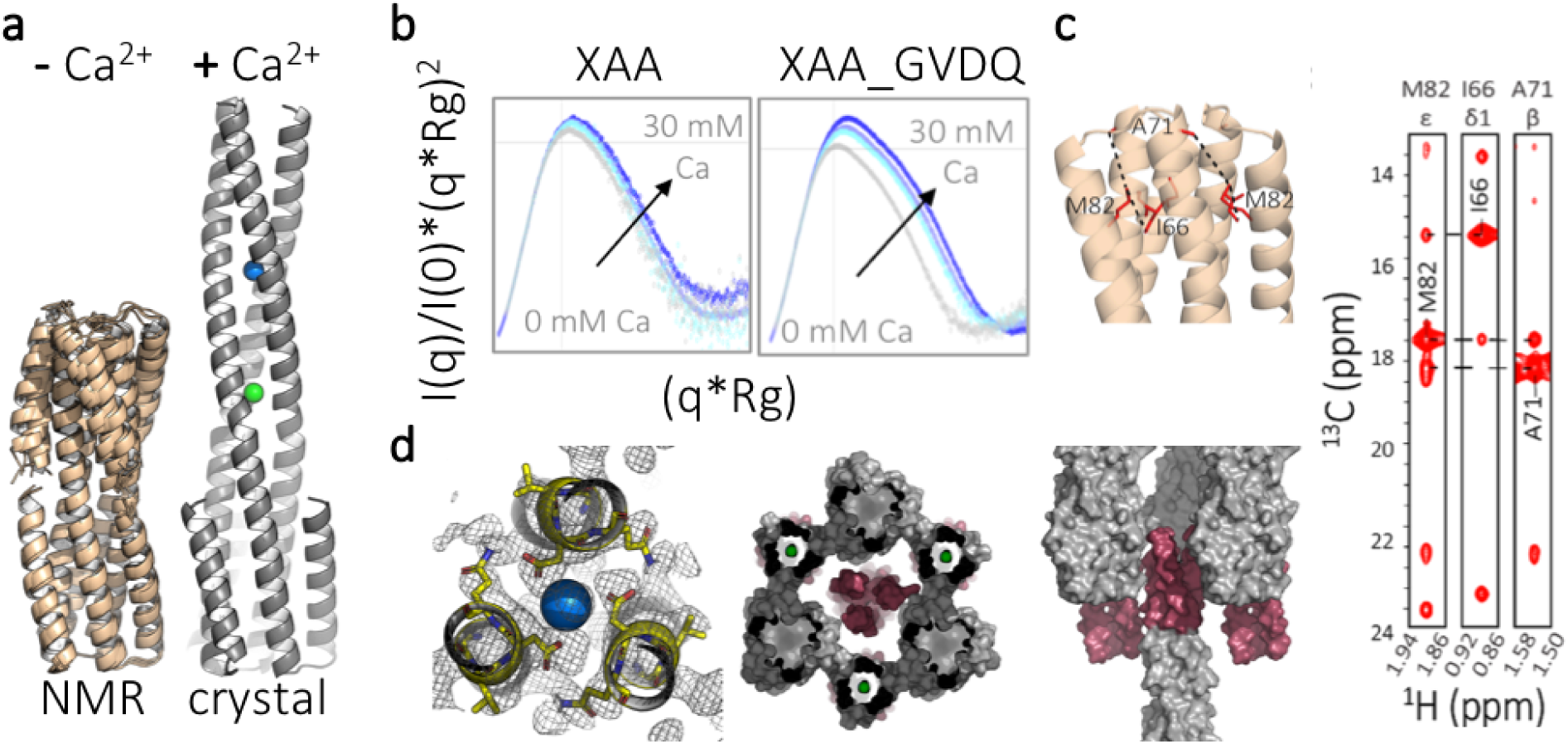
Bistable design XAA_GVDQ is in the short state by NMR and the long state by crystallography. **a**, Summary of crystallographic and NMR characterization of XAA_GVDQ with and without calcium. On the left, Rosetta models based on RDC and chemical shift data shows the protein to be in the short state without calcium (PDB ID 6o0c). On the right, crystal structures shows the protein to be in the long state with calcium (PDB ID 6ny8). **b**, Normalized Kratky plots from SAXS experiments for control protein XAA (linker GGGT) and sample protein XAA_GVDQ measured with up to 30 mM calcium. **c,** 3D C_M_-C_M_H_M_ NOESY strips of XAA_GVDQ indicate the protein is in the short state. **d,** Left panel details crystallographic results at the hinge and hydrogen bond networks with calcium shown as a blue sphere. Middle and right panels show crystal packing from two different views with the flipping helix highlighted in red.

We next sought to determine the structure of the design in the presence of calcium. Overlays of 1D methyl proton spectra show that the protein goes into a non-observable state (several 100s of kDa) in a calcium concentration-dependent manner, even at dilute concentrations and low temperature conditions (Fig. S15). RDCs, amide TROSY spectrum, and 2D 1H-13C HMQC for XAA_GVDQ samples with and without calcium were also recorded (Fig. S15). SAXS analysis of XAA_GVDQ and the control sequence with no aspartate, XAA (linker GGGT), shows calcium dependent changes in the normalized Kratky plot indicating in increase in size in solution (Fig. 4b, Table S4).

Since the calcium-bound state was not observable by NMR, despite efforts to optimize pH, temperature, concentration, calcium, and buffer components, we turned to X-ray crystallography, and sought to determine crystal structures in both the absence and presence of calcium. After extensive screening, we were able to solve a structure in the absence of calcium (Fig. S17, PDB ID 6nxm) in which the flipping helices interact with the inner helices in the same manner as they would in the designed short state, but do so in a domain swapped manner with a symmetry mate to form a hexamer. This is consistent with the observation of the short state by NMR in the absence of calcium; the domain swapping almost certainly arises during crystallization.

We next sought to solve the structure of the design in the presence of calcium. We succeeded in solving the structure in 50 mM calcium acetate by molecular replacement to 2.3 Å resolution (Table S3). The protein is in the designed long state with a metal ion, likely calcium, coordinated by the aspartates in the hinge region (Fig. 4a, PDB ID 6ny8). The ion is most likely calcium given the coordination and the crystallography conditions, which consisted of 0.05 M calcium acetate, 0.1 M sodium acetate pH 4.5, 40% (w/v) 1,2-propanediol at 18°C.

The differences in the crystal and NMR structures of XAA_GVDQ are striking. It is possible that the long state is driven by crystal interaction forces rather than ion coordination, but these crystal specific interactions are likely to be quite small as the flipping helix makes relatively few crystal-lattice interactions (Fig. 4d). On the other hand, since at least >95% of the molecules in solution in the presence of calcium are in the short state by solution NMR, the short state appears to be more favorable in solution. We cannot rule out the possibility that both states are indeed populated in solution, but the barrier to interconversion is high and the long state is removed from solution under NMR conditions: the calcium dependent sample precipitation noted above could reflect the long state being more prone to aggregation, in which case it could be lost in the NMR experiment which is carried out at high protein concentrations.

## Discussion

We have designed a family of de novo proteins in which small changes (3 mutations) in the interface or hinge region can cause the proteins to fold into two distinct and structurally divergent conformations. In contrast to most previous work, the differences in conformation of our designs are relatively large, do not rely on changes in oligomerization state, and both states are well-defined. Most notably, the protein XAA_GVDQ adopts the designed short state by NMR in the absence of calcium, and the designed long state by crystallography in the presence of calcium. Given that changes in three residues in the hinge region can change the conformation of the design (hinge GGHN is long vs. GGGT is short), future work focused on these residues could yield designs that occupy the short and long states in equilibrium and are switchable in solution conditions. Our designed systems, poised as they are between two different states, are excellent starting points for design of triggered conformational changes -- only small or binding free energies would be required to switch conformational states; a particularly interesting possibility would be synthetic environmentally sensitive membrane fusion proteins analogous to those of the viral proteins that inspired the design scheme.

## Materials and Methods

### Synthetic gene construction

Synthetic genes were ordered from Genscript Inc. (Piscataway, NJ, USA) and delivered in pET28b+ *E. coli* expression vectors, inserted at the NdeI and XhoI sites. Genes are encoded in frame with the N-terminal hexahistidine tag and thrombin cleavage site. A stop codon was added at the C-terminus just before XhoI.

### Bacterial protein expression and purification

Plasmids were transformed into chemically competent *E. coli* BL21(DE3)Star (Invitrogen) or Rosetta(DE3)pLysS (QB3 MacroLabs). Starter cultures were grown in Lysogeny Broth (LB) media with 50 μg/mL kanamycin overnight with shaking at 37°C. 5 mL of starter culture was used to inoculate 500 mL of 2xTY with 50-100 μg/mL kanamycin at 37°C until OD600 = 0.6. Cultures were cooled to 18°C and induced with 0.1mM IPTG at OD600 = 0.8 and grown overnight. Cells were harvested by centrifugation at 3500 rcf for 15 minutes at 4°C and resuspended in lysis buffer (25 mM Tris pH 8.0, 300 mM NaCl, 20 mM imidazole), then lysed by sonication. Lysates were cleared by centrifugation at 14,500 rcf for at least 45 minutes at 4°C. Supernatant was applied to 0.5-1 mL Ni-NTA resin (Qiagen) in gravity columns pre-equilibrated in lysis buffer. The column was washed thrice with 10 column volumes (CV) of wash buffer (25 mM Tris pH 8.0, 300 mM NaCl, 20 mM imidazole). Protein was eluted with 25 mM Tris pH 8.0, 300 mM NaCl, 250 mM imidazole, 0.5 mM TCEP. Proteins were buffer exchanged using FPLC (Akta Pure, GE) and a HiPrep 26/10 Desalting column (GE) into 25 mM Tris pH 8.0, 300 mM NaCl. The hexahistidine tag was removed by thrombin cleavage by incubating with 1:5000 thrombin (EMD Millipore) overnight at room temperature. Proteins were further purified by gel filtration using FPLC and a Superdex 75 10/300 GL (GE) size exclusion column.

### Circular dichroism (CD) measurements

CD wavelength scans (260 to 195 nm) and temperature melts (25°C to 95°C) were measured using an AVIV model 410 CD spectrometer. Temperature melts monitored absorption signal at 222 nm and were carried out at a heating rate of 4°C/min. Protein samples were prepared at 0.25 mg/mL in PBS in a 1 mm cuvette.

### Small angle X-ray scattering (SAXS) measurements

Proteins were purified by gel filtration using FPLC (Akta Pure, GE) and a Superdex 75 10/300 GL (GE) into SAXS buffer (20 mM Tris pH 8.0, 150 mM NaCl, and 2% glycerol). Fractions containing trimers were collected and concentrated using Spin-X UF 500 10 MWCO spin columns (Corning) and the flow through from the spin columns was used as buffer blanks. Samples were prepared at concentrations of 3-8 mg/ml. SAXS measurements were made at SIBYLS 12.3.1 beamline at the Advanced Light Source (ALS). The light path is generated by a super-bend magnet to provide a 1012 photons/sec flux (1 Å wavelength) and detected on a Pilatus3 2M pixel array detector. Each sample is collected multiple times with the same exposure length, generally every 0.3 seconds for a total of 10 seconds resulting in 30 frames per sample.

### Protein crystallography, x-ray data collection, and structure determination

Proteins were purified by gel filtration using FPLC (Akta Pure, GE) and a Superdex 75 10/300 GL (GE) into 20 mM Tris pH 8.0, 150 mM NaCl and concentrated to 3-10 mg/ml, most commonly 8 mg/ml. Samples were screened using a Mosquito (TTP Labtech) with JCSG I-IV (Qiagen), Index (Hampton Research), and Morpheus (Molecular Dimensions) screens. Sitting drops of 1:1 mixture of 150-200 nL protein and 150-200 nL reservoir solution at 4°C or 18°C. Crystallization conditions are listed in Table S2. The X-ray data sets were collected at the Berkeley Center for Structural Biology beamlines 8.2.1 and 8.3.1 of the Advanced Light Source at the Lawrence Berkeley National Laboratory (LBNL). Data sets were indexed and scaled using HKL2000 (Otwinowski and Minor, 1997) or XDS (Kabsch, 2010). Structures were determined by the molecular-replacement method with the program PHASER (McCoy et al., 2007) within the Phenix suite(Adams et al., 2010) using the Rosetta design models as for the initial search model. The molecular replacement results were used to build the design structures and initiate crystallographic refinement and model rebuilding. Structure refinement was performed using the phenix.refine program (Adams et al., 2010; Afonine et al., 2010). Manual rebuilding using COOT (Emsley and Cowtan, 2004) and the addition of water molecules allowed construction of the final models. Summaries of statistics are provided in Table S3.

### NMR spectroscopy and structure determination

All experiments were recorded at 37 °C on an 800 MHz Bruker Avance III spectrometer equipped with a TCI cryoprobe. NMR samples contained 20 mM Sodium Phosphate pH 6.2, 100mM NaCl, 0.01% NaN_3_, 1 U Roche protease inhibitor cocktail in 95% H_2_O/5% D_2_O. For backbone and methyl assignments, labeled samples were prepared using well-established protocols (Tugarinov et al., 2006; Tugarinov and Kay, 2003). Briefly, backbone assignments were performed on uniformly ^13^C/^15^N, perdeuterated protein samples using TROSY-based 3D experiments, HNCO, HNCA and HN(CA)CB. Side-chain methyl assignments were performed starting from the backbone assignments and using unambiguous NOE connectivities observed in 3D C_M_-C_M_H_M_, 3D C_M_-NH_N_ and 3D H_N_H_α_-C_M_H_M_ SOFAST NOESY experiments, recorded on a U-[^15^N, ^12^C, ^2^H] AI(LV)^proS^ (Ala ^13^Cβ, Ile ^13^Cδ1, Leu ^12^Cδ1/^13^Cδ2, Val ^12^Cγ1/^13^Cγ2) selective methyl-labeled sample. The calcium titrations for XAA_GVDQ were performed in 20 mM Tris-HCl pH 7.2, 100 mM NaCl and 0, 0.5, 1, 2, 5, 10, 15, 20, 25, 30 and 50 mM CaCl_2_ buffer conditions.

Backbone amide ^1^D_NH_ residual dipolar couplings (RDCs) were obtained using 2D ^1^H-^15^N transverse relaxation-optimized spectroscopy (ARTSY) (Fitzkee and Bax, 2010). For XAA_GVDQ sample, the alignment media was 10 mg/mL Pf1 phage (ASLA Biotech) for experiments performed without calcium and 15 mg/mL for experiments performed in the presence of 5 mM CaCl_2_. The ^2^H quadrupolar splittings were 8.7 Hz and 9.3 Hz in the absence and presence of CaCl_2_, respectively. For XAA sample, the alignment media was 10 mg/mL Pf1 phage (ASLA Biotech) and the ^2^H quadrupolar splitting was equal to 6.2 Hz. All NMR data were processed with NMRPipe(Delaglio et al., 1995) and analyzed using NMRFAM-SPARKY(Lee et al., 2015). The Chemical Shift Deviations (CSD) of ^13^C methyls (in p.p.m.) were calculated using the equation 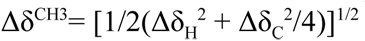 and of ^15^N amides (in p.p.m.) using the equation 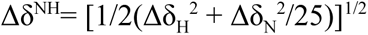.

Structure modeling from our NMR data was performed using Rosetta’s fold and dock protocol(Das et al., 2009; DiMaio et al., 2011; Sgourakis et al., 2011; Tyka et al., 2011). Two sets of calculations were performed, using different input data (i) backbone chemical shifts and RDCs (to model the structure of XAA_GVDQ in the presence and absence of calcium, Fig 4a bottom) and (ii) backbone chemical shifts, long-range NOEs and RDCs data (to calculate the *de novo* NMR structures for XAA (Fig. 2e) and XAA_GVDQ (Fig. 4d))(Nerli et al., 2018). In (ii), we used a uniform upper distance bound of 10 Å for all 92 NOE restraints for XAA and for all 139 NOE restraints for XAA_GVDQ (Table S5). The top 10 lowest-energy structures from each calculation showing good agreement with RDCs and chemical shifts and a minimum number of NOE violations were deposited in the PDB, under accession code 6O0I for XAA (BMRB ID 30574) and 6O0C for XAA_GVDQ mutant M4L (BMRB ID 30573).

## Acknowledgements

We thank X. Ren, J. Hurley, G. Meigs, P. Adams, B. Sankaran, and A. Kang for help with X-ray crystallography. We thank K. Burnettt, G. Hura, and J. Tanamachi for help with SAXS. We thank C. Nixon and S. Marqusee for help with CD. We thank B. Belardi, M. Vahey, S. Son, S. Ovchinnikov, and the rest of the Fletcher and Baker laboratories for discussion. We thank UW Hyak supercomputer and Rosetta@Home volunteers (https://boinc.bakerlab.org) for computing resources.

## Author contributions

K.Y.W., S.E.B., P.-S.H., N.G.S and D.B., designed the research.

K.Y.W. designed, performed, and analyzed data for CD, SEC-MALS, SAXS, and X-ray crystallography experiments.

D.M., A.C.M. and N.G.S. performed NMR experiments and data analysis.

M.J.B. designed and analyzed data for X-ray crystallography experiments.

S.N. performed NMR structure calculations.

L.C. and D.M. performed SEC-MALS experiments and NMR sample preparation.

All authors contributed to experimental design, discussed results, and commented on the manuscript.

## Supplementary Materials

### Supplemental Figures

**Fig S1.**
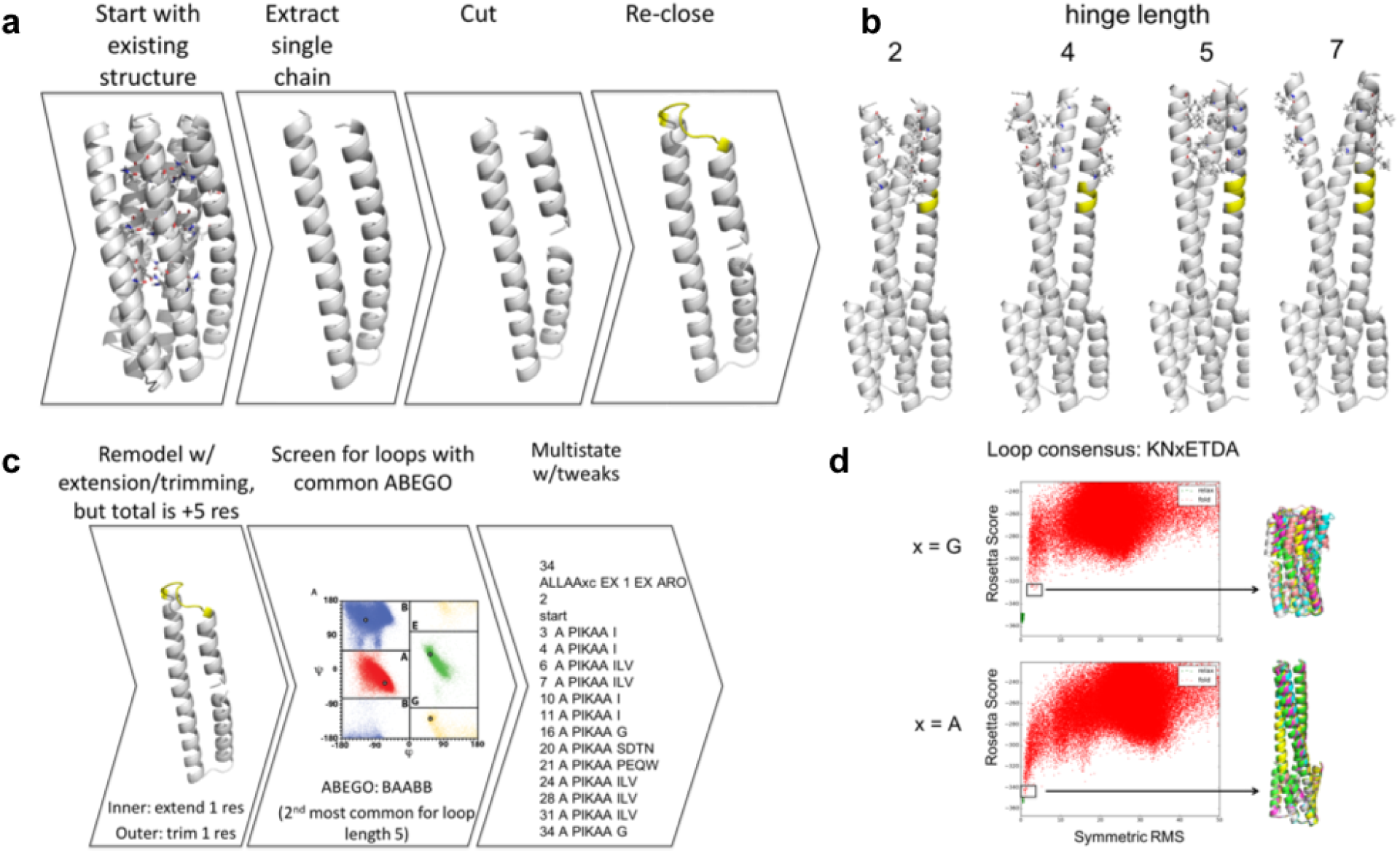
Round 1 design scheme. **a,** Starting with previously characterized design 2L6HC3_13 variant (i.e., 2L6HC3_XAAA)^1^, the protein is rearranged to form the desired architecture. Helix capping residues were incorporated at the N-terminus, and hydrogen bond networks within the flipping helix were redesigned using Rosetta to hydrophobic residues for stability. **b,** Different hinge lengths were tested for ability to pack in the long state. **c**, Loop closure using the determined hinge length. **d**, Testing for sequences that are predicted by Rosetta to fold in the short or long state.

**Fig S2.**
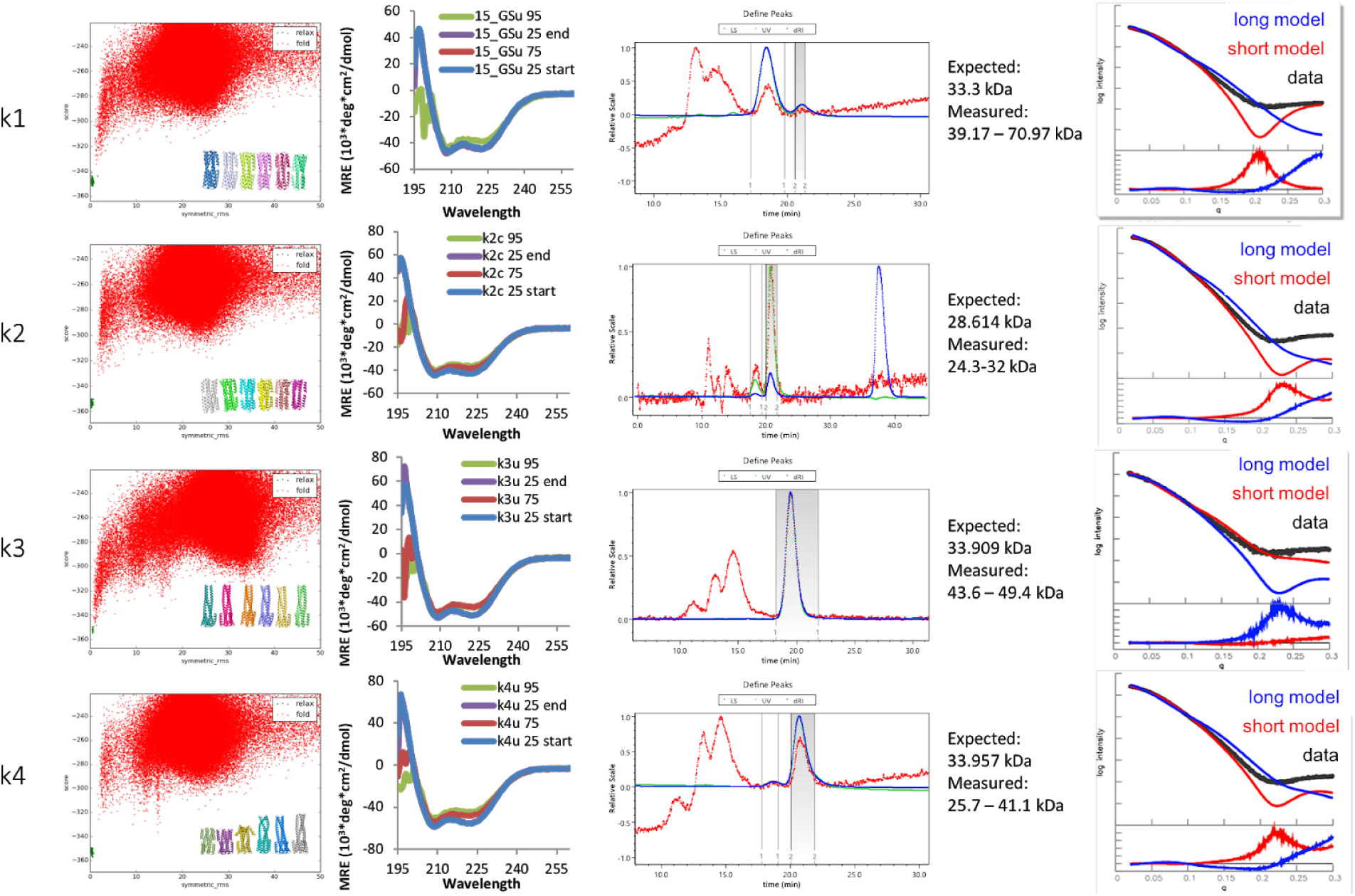
Characterization of round 1 designs. (first column) Rosetta folding predictions, (second column) circular dichroism (CD) measurements, (third column) size exclusion chromatography with multi-angle light scattering (SEC-MALS) measurements, and (fourth column) small angle x-ray scattering (SAXS) measurements of designs k1-k4.

**Fig S3.**
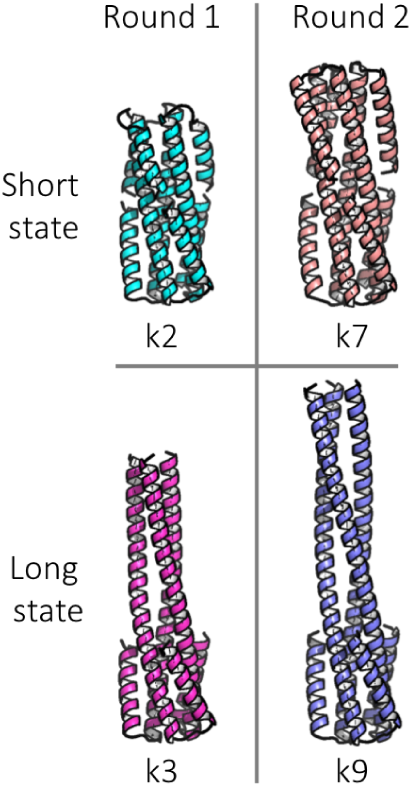
Comparison of round 2 to round 1 designs. Round 2 designs have longer flipping helices, compared to round 1 designs, to facilitate discrimination by SAXS.

**Fig S4.**
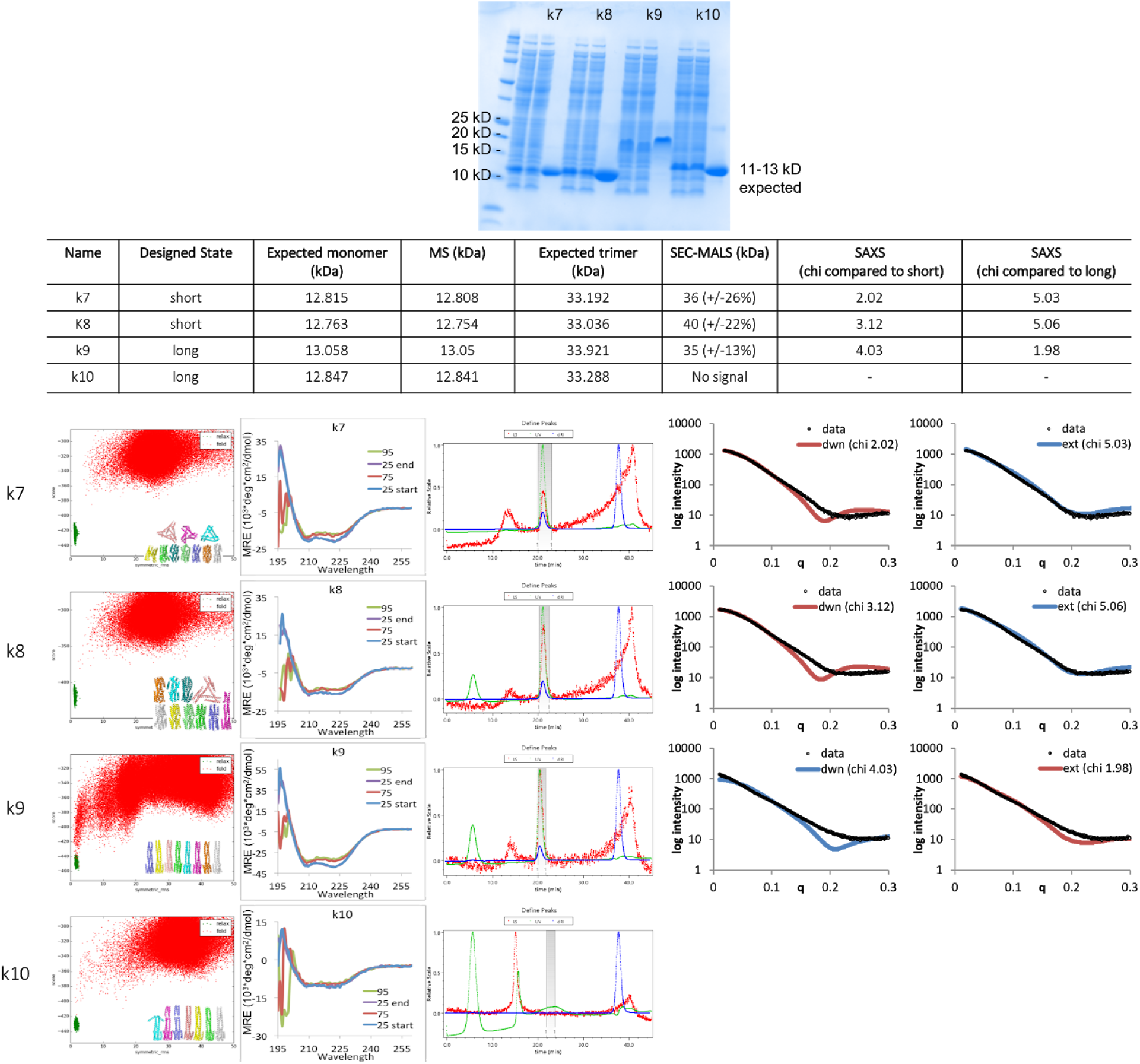
Characterization of round 2 designs. (top) SDS-PAGE gel, (second row) table summary of characterization results, (first column) Rosetta folding predictions, (second column) CD measurements, (third column) SEC-MALS measurements, and (fourth and fifth column) SAXS measurements of designs k7-10. Design k9 runs larger than expected on SDS-PAGE gels, but this can sometimes occur for highly helical proteins.

**Fig S5.**
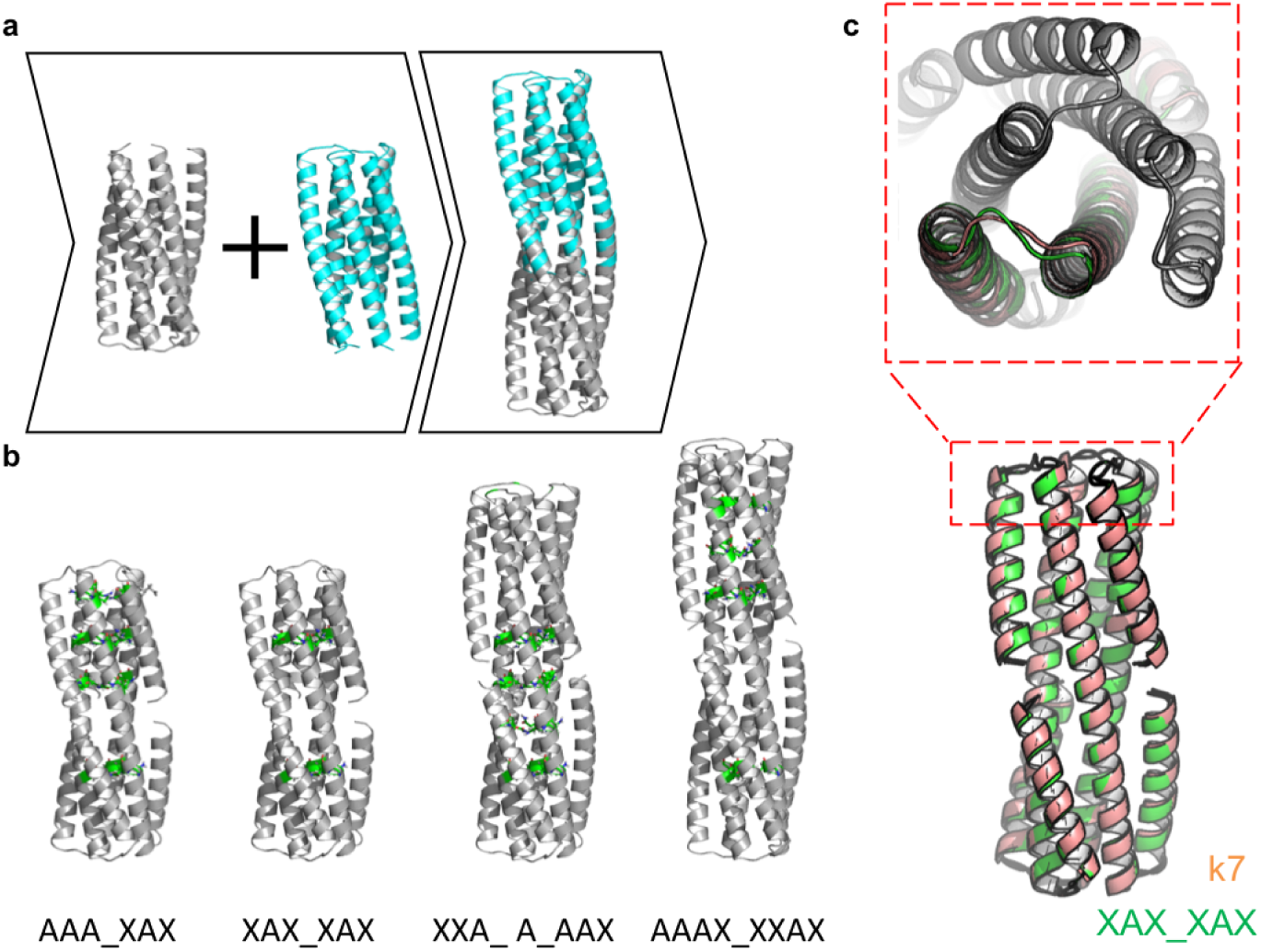
Round 3 design scheme and comparison to round 2. **a,** Backbone construction by fusing 2L6HC3_13 variant (2L6HC3_XAAA) (grey) to 2L6HC3_23 (cyan)^1^ via the inner three helices. Outer helices are trimmed to remove clashes and the exposed inner helices were redesigned using Rosetta to remove surface exposed hydrogen bond networks. **b**, Four variants with different fusion positions, placement of hydrogen bond networks (highlighted in green), and loop connectivity. **c,** Compared to the second generation designs (k7, orange), the flipping helix of round 3 design (XAX_XAX, green) is positioned slightly closer to the inner helices in the short state and the termini of the hinge are slightly closer together.

**Fig S6.**
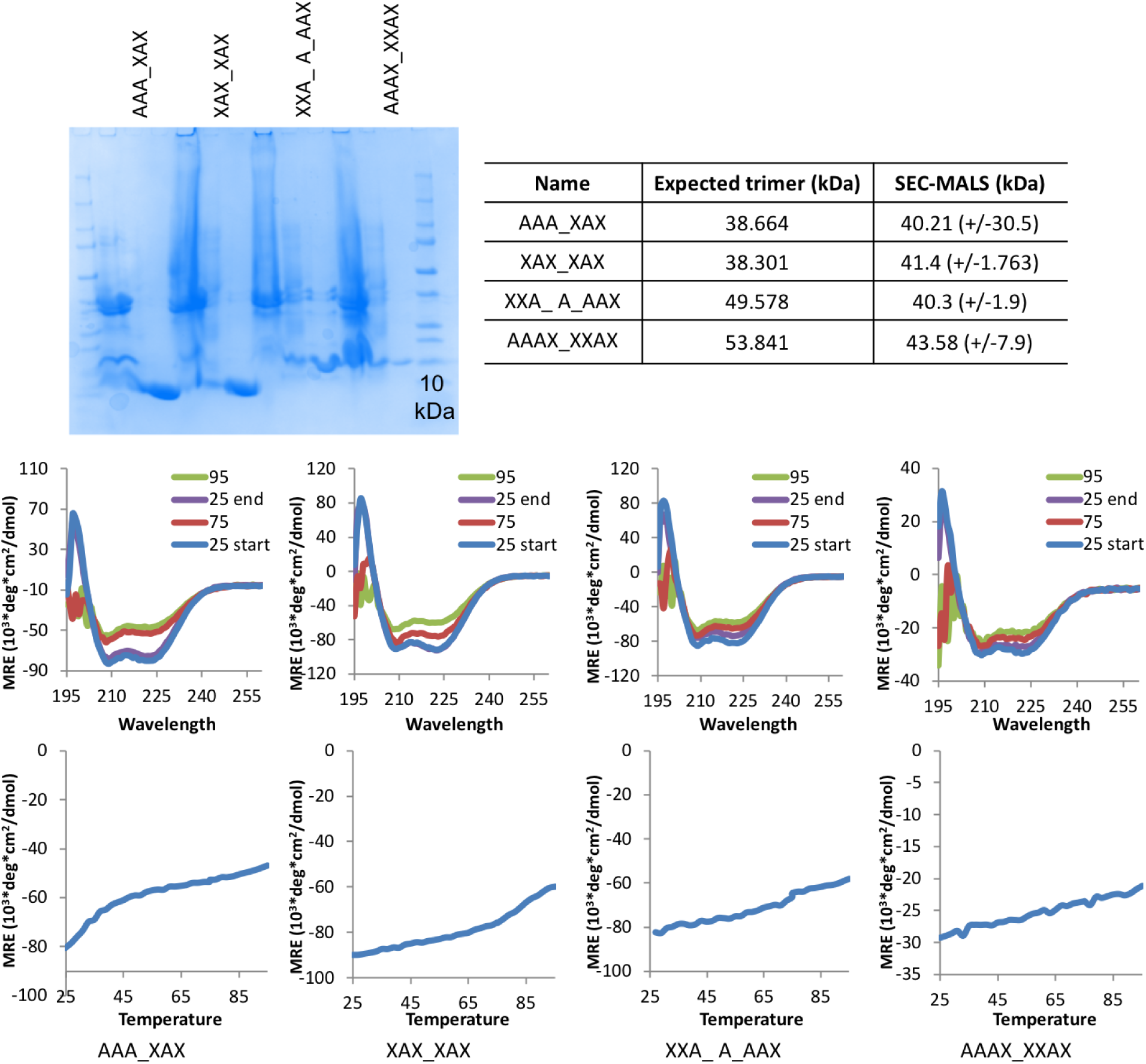
Characterization of round 3 designs. (top) SDS-PAGE gel and table summary of characterization results, and (bottom) CD measurements for round 3 designs. Top row shows wavelength scans at 25°C, 75°C, 95°C, and samples cooled back to 25°C. Bottom row shows temperature melts of constructs from 25°C to 95°C.. AAA_XAX is also referred to as AAA. Both XXA_A_AAX and AAAX_XXAX appear smaller than would be expected of trimers by SEC-MALS.

**Fig S7.**
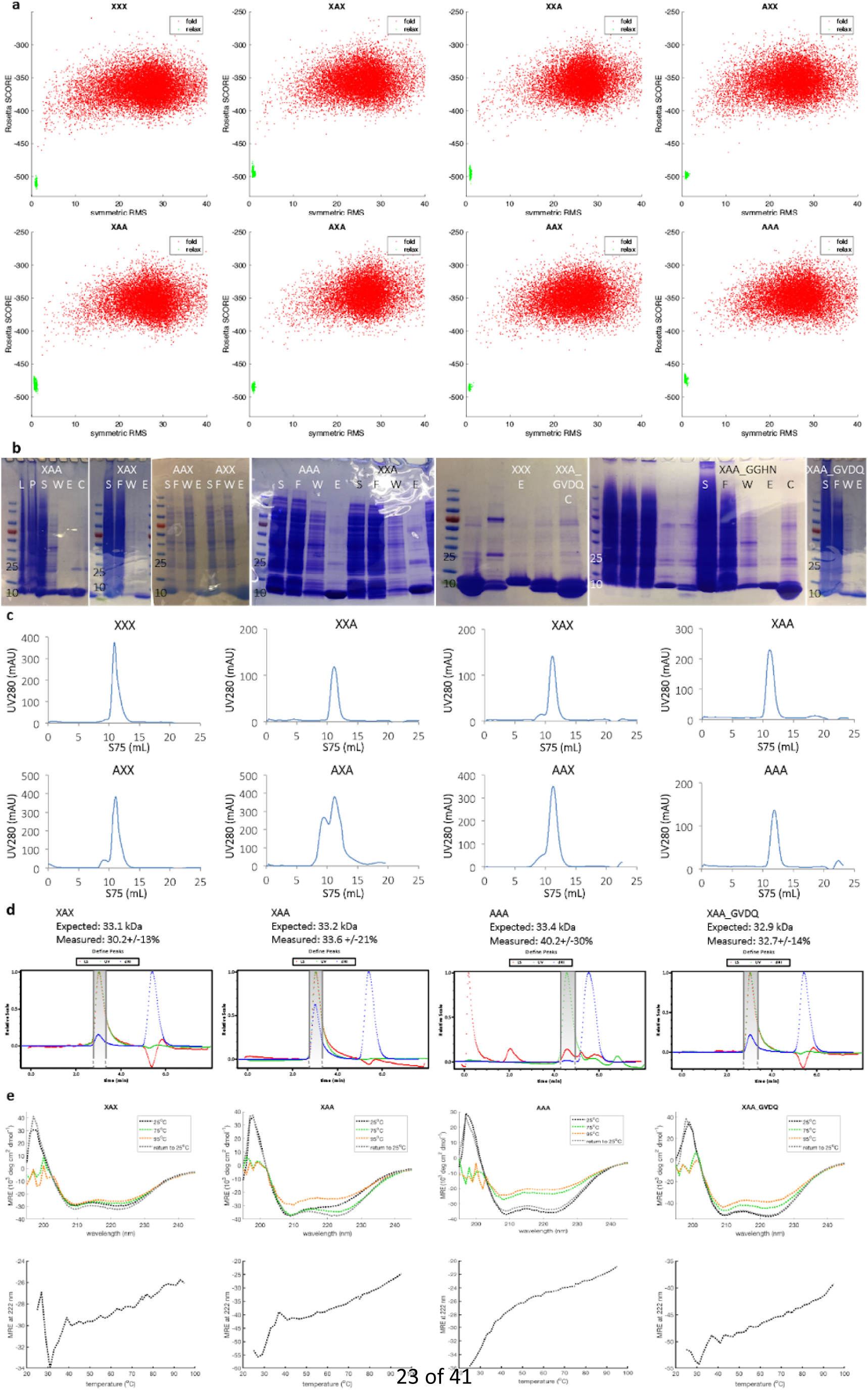
Protein expression and nickel purification. **a,** Rosetta folding predictions plotted with Rosetta energy on the y-axis and symmetric RMS to input model on the x-axis. Green points are relaxation runs on the model (of the short state only) and red points are individual folding trajectories (based on sequence alone). **b,** SDS-PAGE gel showing various steps during nickel purification to verify expression and purity of designs. L = cell lysate; P = pellet; S = supernatant; F = flow through; W = wash 1 (of 3); E = elution; C = thrombin cleaved. **c,** FPLC purification. Absorbance at 280 nm vs. column volume for protein samples run on a Superdex 75 10/300 GL column in 20 mM Tris, pH 8, 150 mM NaCl, 2% glycerol. Most samples were monodisperse or have a relatively small aggregation peak. AXA has a large soluble aggregate peak, but has a prominent peak at the expected position for a trimer. **d,** SEC-MALS measurements. Expected mass is calculated for trimers after thrombin cleavage to remove histidine purification tag. **e,** CD measurements. (top) wavelength scans at 25°C, 75°C, 95°C, and samples cooled back to 25°C and (bottom) temperature melts of constructs from 25°C to 95°C.

**Fig S8.**
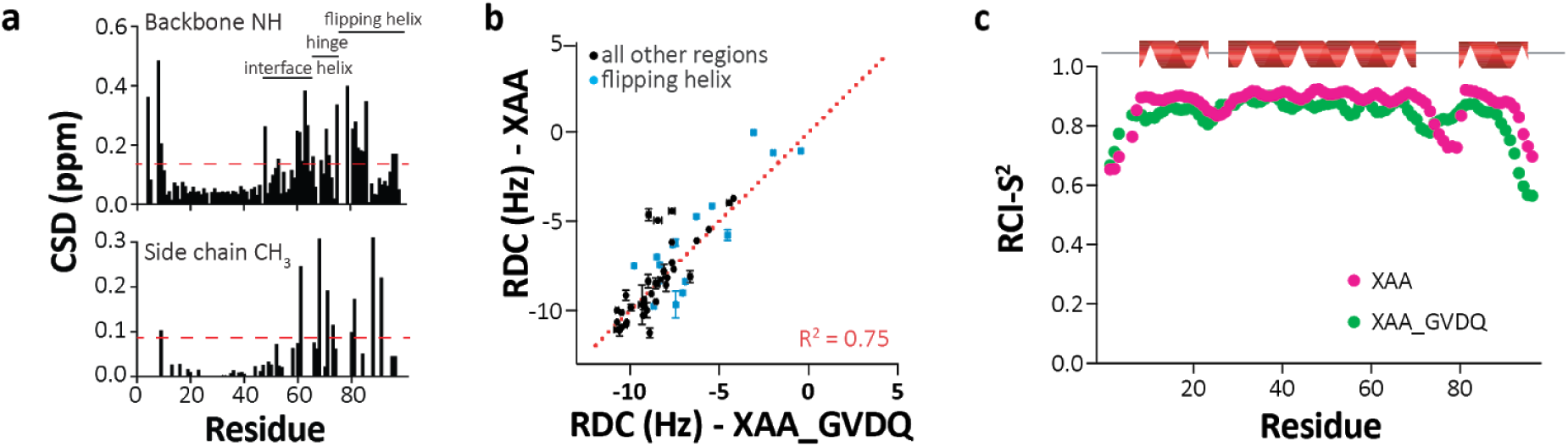
Detailed characterization of XAA. **a,** Chemical shift deviations (CSDs) for amides and methyls shared between XAA and XAA_GVDQ show differences in the flipping helix interface to the inner helices. **b,** RDC experiments for XAA vs XAA_GVDQ. **c,** Estimated backbone order parameter (Random Coil Index S^2^) derived from the chemical shifts^3^ is shown for XAA and XAA_GVDQ residues. Lower RCI-S^2^ indicates flexibility, higher RCI-S^2^ indicates rigidity. Secondary Structure Index (SSI), predicting three helical segments here, are derived from TALOS-N analysis of backbone (^13^C^a^, ^13^C’, ^15^N, ^1^H^a^ and ^1^H^N^) and ^13^C^b^ chemical shifts.

**Fig S9.**
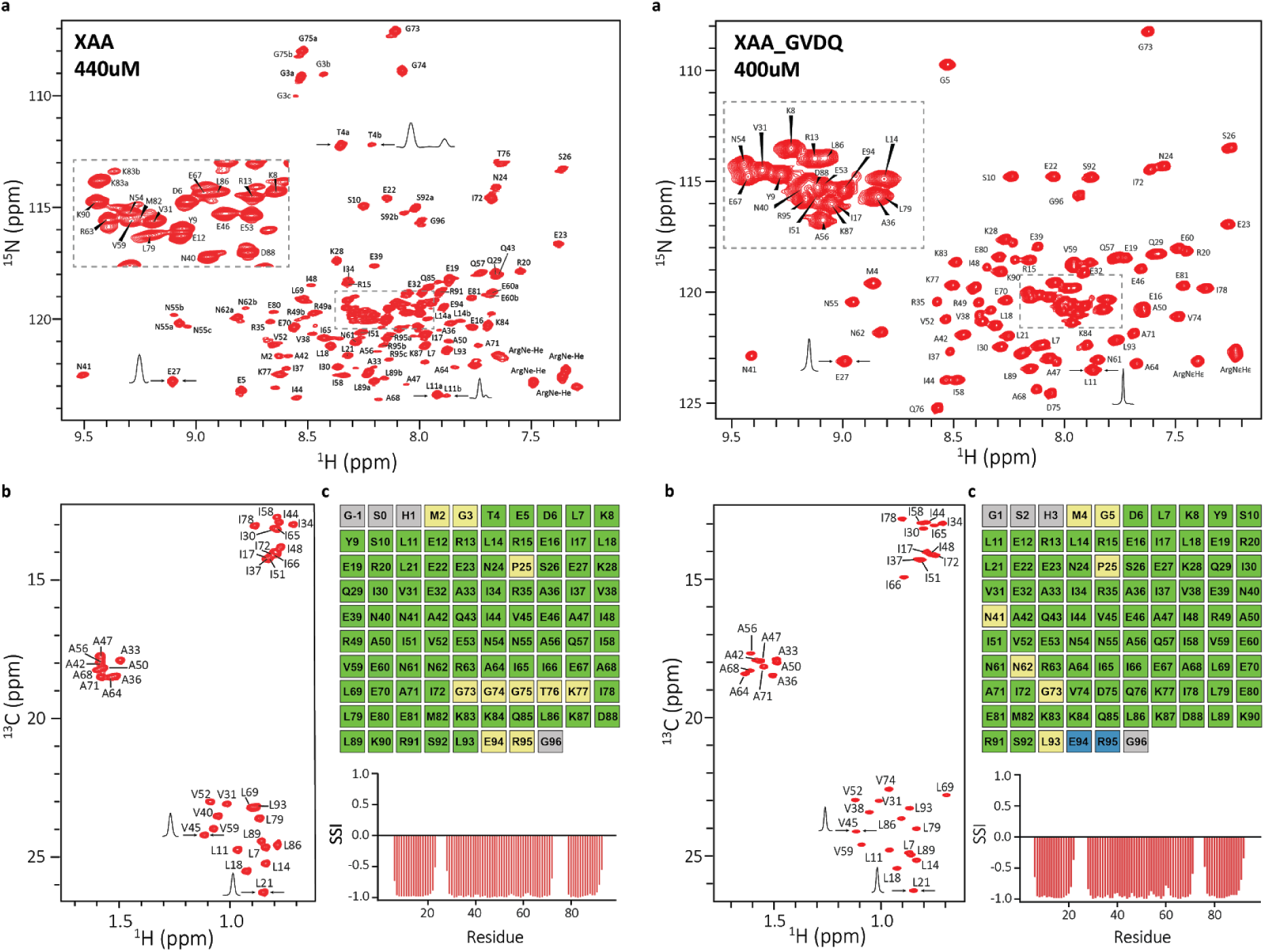
Backbodne and methyl NMR assignments for XAA (left) and XAA_GVDQ (right) **a**, Fully assigned 2D ^1^H-^15^N TROSY-HSQC spectrum. 1D slices are shown with arrows in both spectra to indicate the line broadening on XAA and the presence of a minor conformation in various residues distributed to the entire structure of XAA which turns to be a heterogeneity. **b,** Fully assigned 2D ^1^H-^13^C HMQC for AI(LV)^proS^ samples. **c,** (top) TALOS-N validation data for backbone assignments. Assignment validation, green: highly confident assignments. yellow: acceptable within limits. blue: dynamic residues. grey: no classification. (bottom) Secondary Structure Index (SSI) derived from TALOS-N analysis of backbone (^13^C_a_, ^13^C’, ^15^N, ^1^H_a_ and ^1^H_N_) and ^13^C_b_ chemical shifts. Positive SSI values are consistent with β-strand and negative values are consistent with α helical structure.

**Fig S10.**
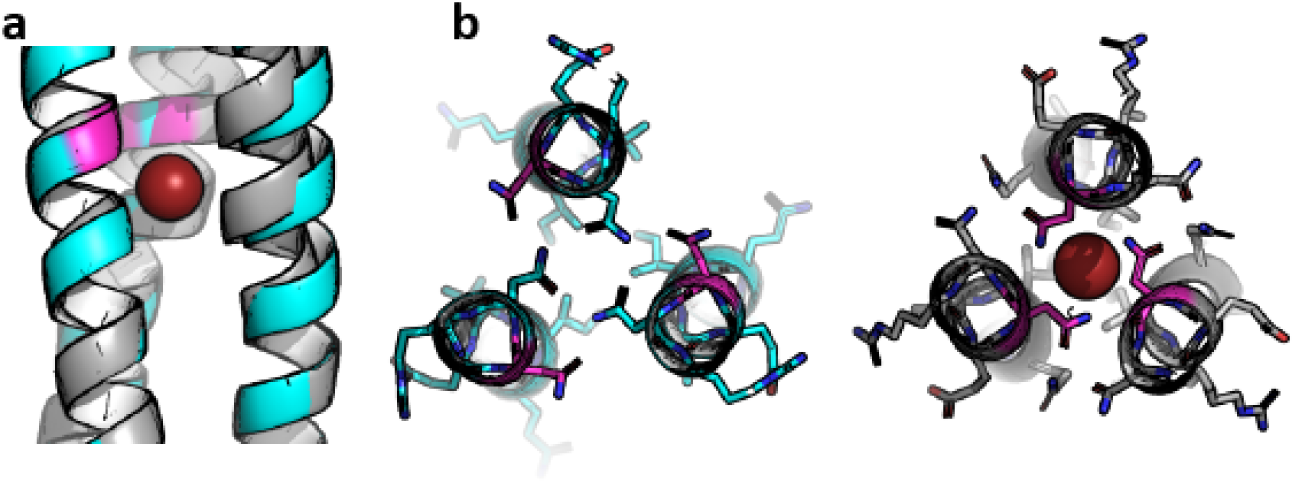
Bromine coordination site in AAA. Model (cyan) and crystal (grey) crystal structure of AAA is shown near N61 (magenta), where there is deviation between the model (left) and the crystal due to the presence of bromine (right).

**Fig S11.**
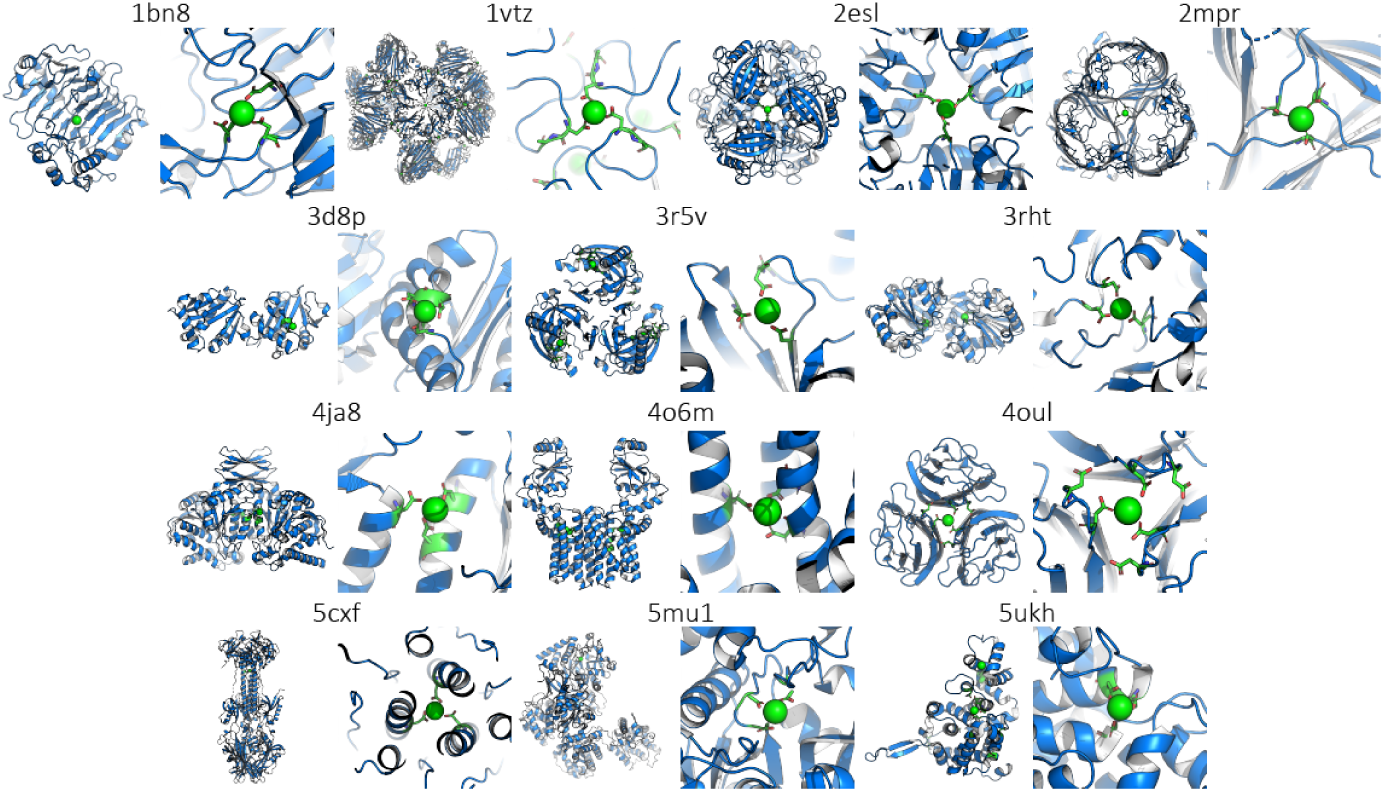
Calcium binding by three asparatates from the PDB. The MetalPDB database (http://metalweb.cerm.unifi.it/search/metal/) was used to curate 13 structures with calcium coordinated by three aspartic acids (of 4520 labeled as calcium binding). For each structure, the PDB ID is given at the top, the full structure shown on the left, and a close up of the binding site shown on the right. Calcium and coordinating aspartates are shown in green.

**Fig S12.**
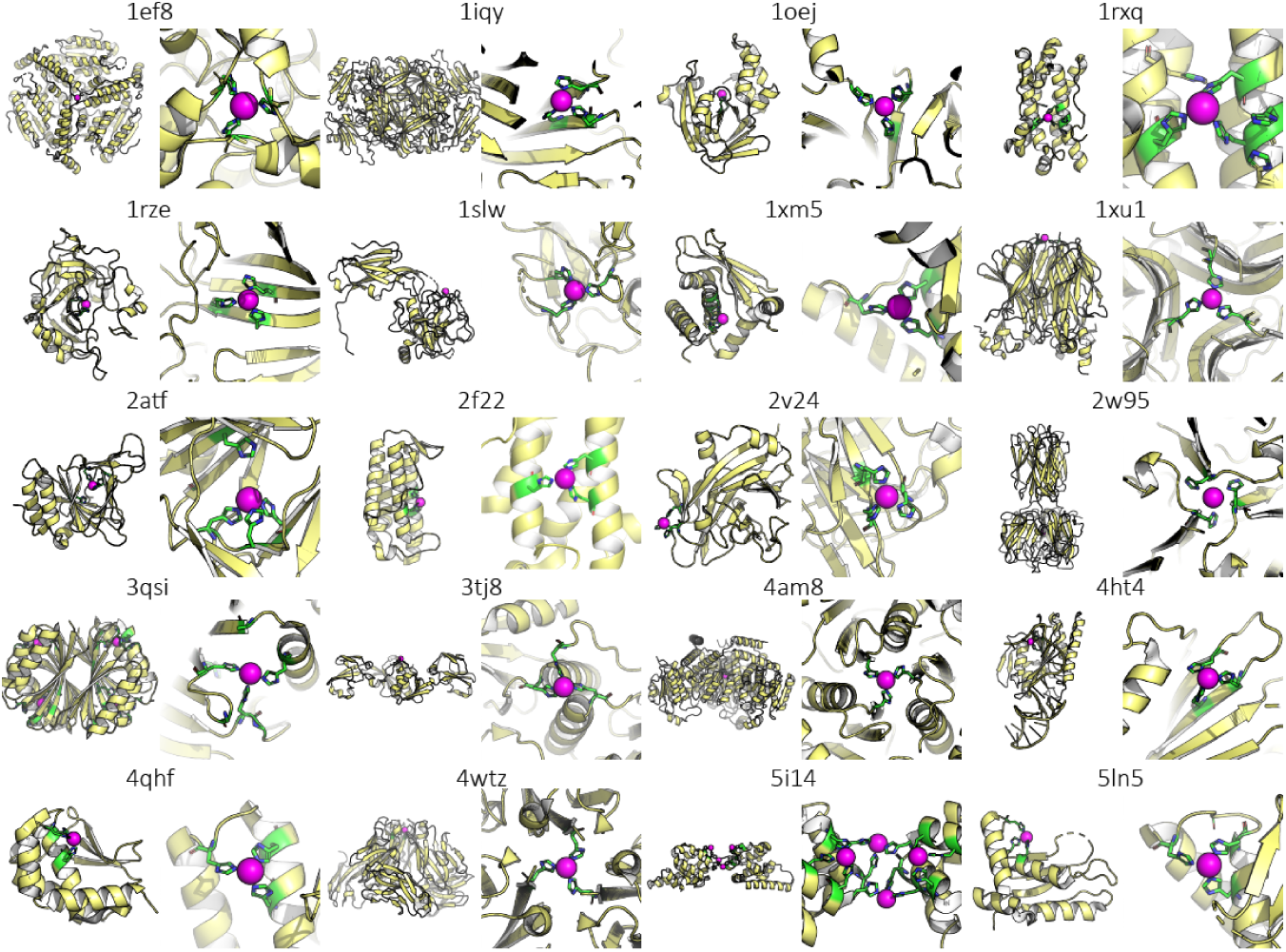
Nickel binding by three histidines from the PDB. The MetalPDB database (http://metalweb.cerm.unifi.it/search/metal/) was used to curate 20 structures with zinc coordinated by three histidines (of 861 labeled as nickel binding). For each structure, the PDB ID is given at the top, the full structure shown on the left, and a close up of the binding site shown on the right. Nickel is shown in magenta and coordinating histidines are shown in green.

**Fig S13.**
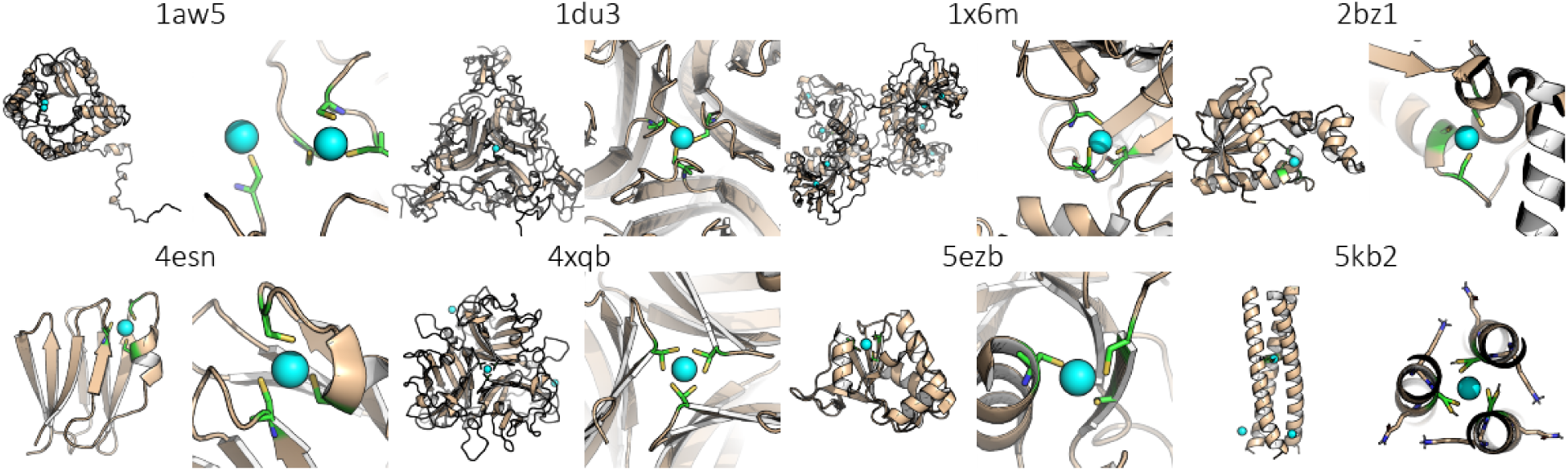
Zinc binding by three cysteines from the PDB. The MetalPDB database (http://metalweb.cerm.unifi.it/search/metal/) was used to curate 8 structures with zinc coordinated by three cysteines (of 5467 labeled as zinc binding). For each structure, the PDB ID is given at the top, the full structure shown on the left, and a close up of the binding site shown on the right. Zinc is shown in cyan and coordinating cysteines are shown in green.

**Fig S14.**
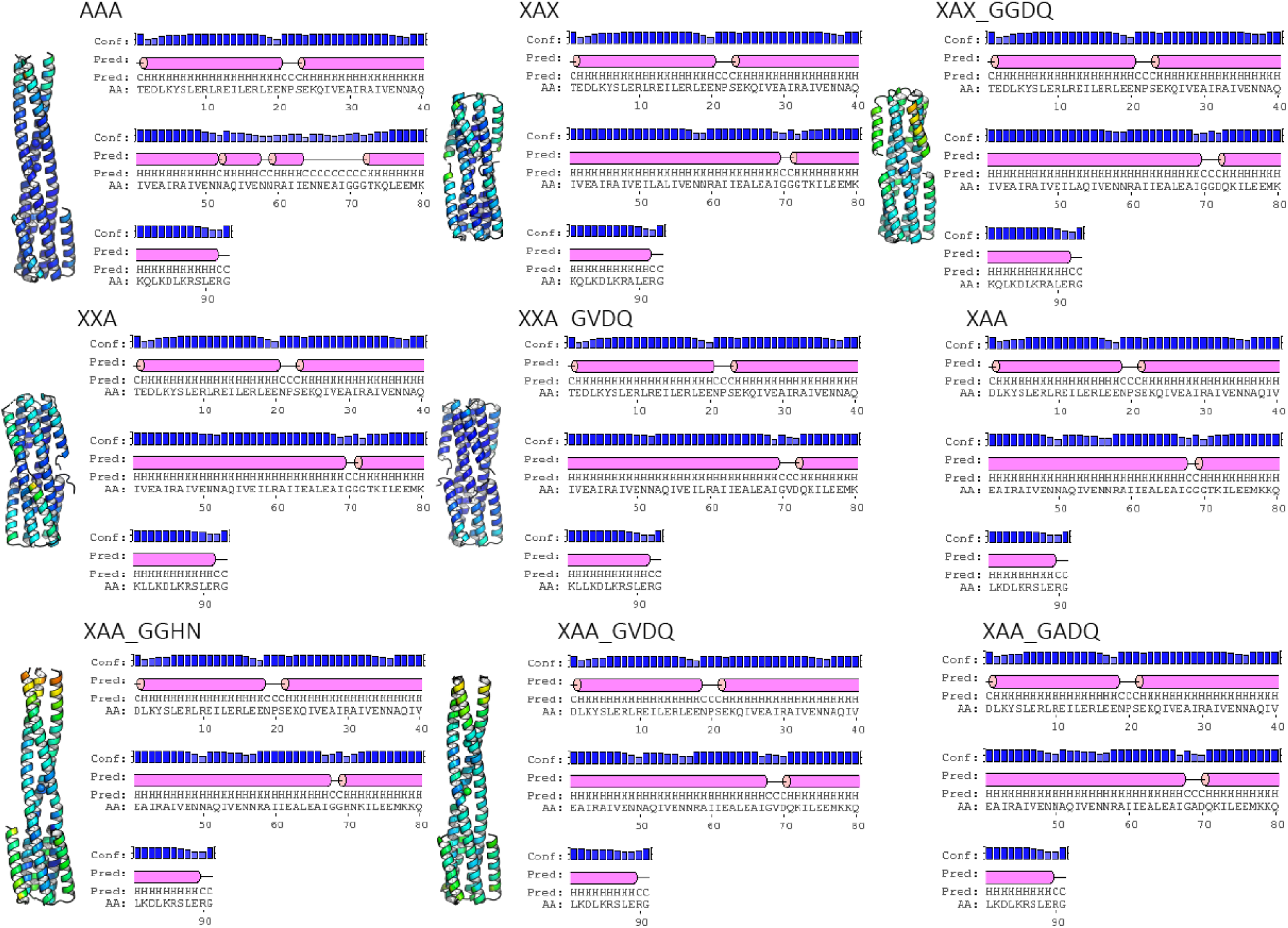
B-factors and helical propensities of selected designs. B-factors are show on ribbon diagrams by color (blue is low, red is high; not normalized across structures). PSIPRED predictions are shown diagrammatically and by letter designation^2^.

**Fig S15.**
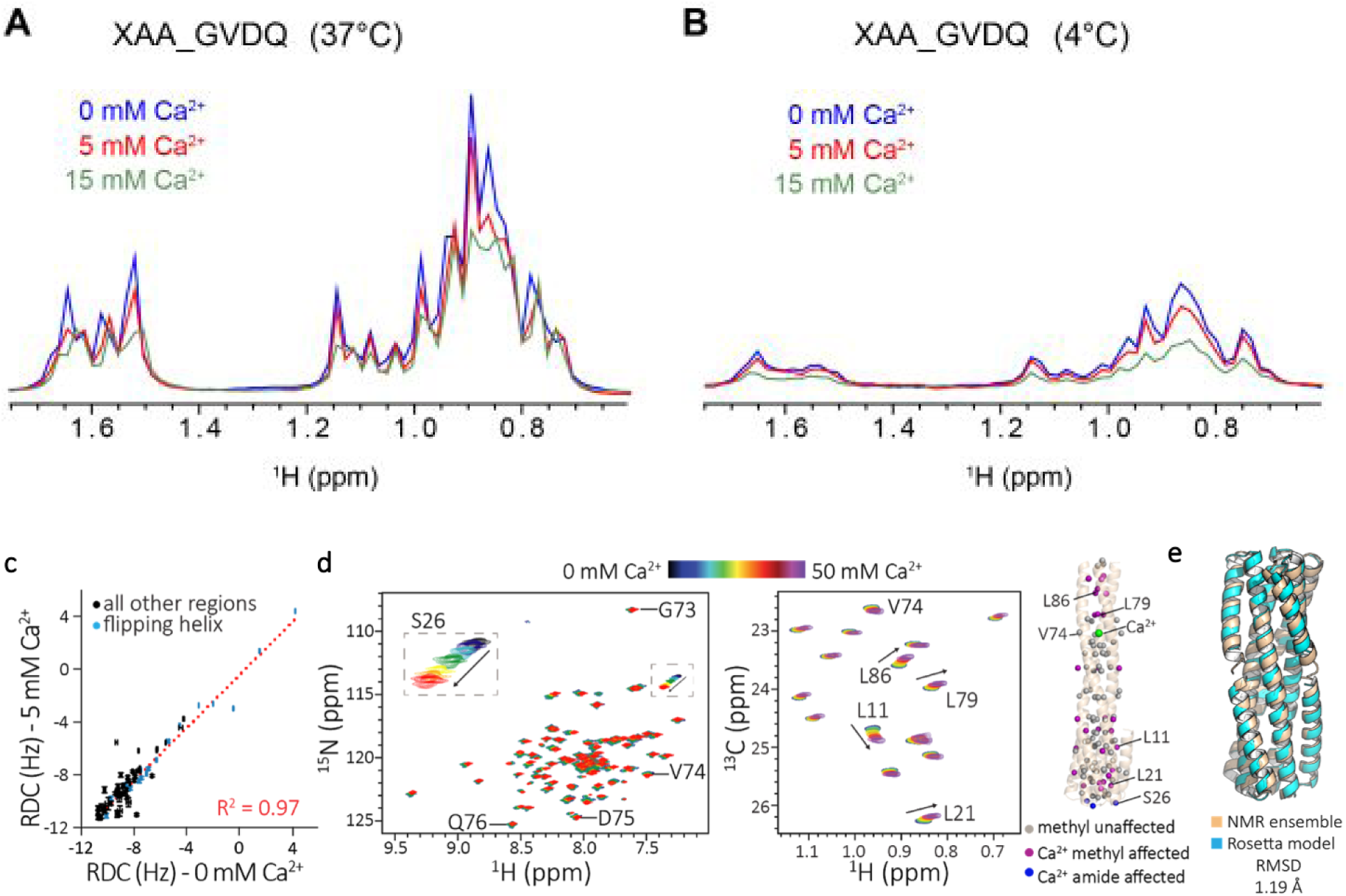
1D methyl proton spectra for XAA_GVDQ with calcium. Comparison of 1D projections along the ^1^H dimension extracted from 2D ^1^H-^13^C methyl SOFAST HMQC experiments recorded at **a,** 37 °C or **b,** 4°C at a ^1^H field strength of 800 MHz. For experiments at 37 °C, acquisition parameters were 4 scans/FID with a recycle delay of 0.2 sec and 128 / 1024 complex points in the ^13^C / ^1^H dimensions. For experiments at 4 °C, acquisition parameters were 8 scans/FID with a recycle delay of 0.2 sec and 128 / 1024 complex points in the ^13^C / ^1^H dimensions. Each experiment was recorded in the presence of varying concentrations of calcium using 105 μM MAI(LV)^proS^ labeled XAA_GVDQ in NMR buffer (100 mM NaCl, 20 mM Tris pH 7.2, 2% (v/v) d_8_-glycerol). **c,** RDC experiments with 0 mM or 5 mM calcium. Amide groups corresponding to the C-terminal helix are colored blue. **d,** Calcium titration of XAA_GVDQ construct. (left) Overlay of 2D ^1^H-^15^N TROSY-HSQC spectra and (center) 2D ^1^H-^13^C HMQC spectra of AI(LV)^proS^ sample in different calcium concentrations (0-50 mM). Non-specific calcium binding is observed. G73, V74, D75 and Q76 are the residues expected in calcium binding and S26, L11, L21, L79 and L89 are the residues observed. (right) Distribution of calcium unaffected methyl (grey), methyl affected (magenta), and calcium amide affected (blue) residues overlayed on the long conformation expected from crystallographic data. **e**, De novo NMR structure for XAA_GVDQ (beige, PDB ID 6o0c) compared to design model (cyan) confirms the protein is in the compact conformation (RMSD = 1.19 Å).

**Fig S16.**
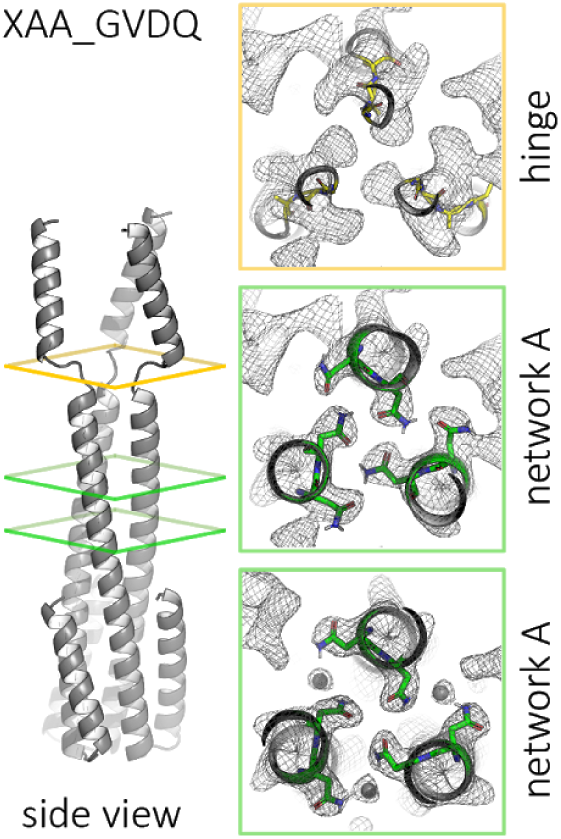
Crystal structure of XAA_GVDQ without calcium (PDB ID 6nxm) The protein crystallizes as a domain swapped hexamer (only trimer shown) with contacts in the flipping helix replicating the contacts of the short model state, but with another molecule of XAA_GVDQ rather than with self. Details showing the GVDQ hinge region forms a very flexible linker that allows the domain swap. Designed hydrogen bond networks remain intact.

**Fig S17.**
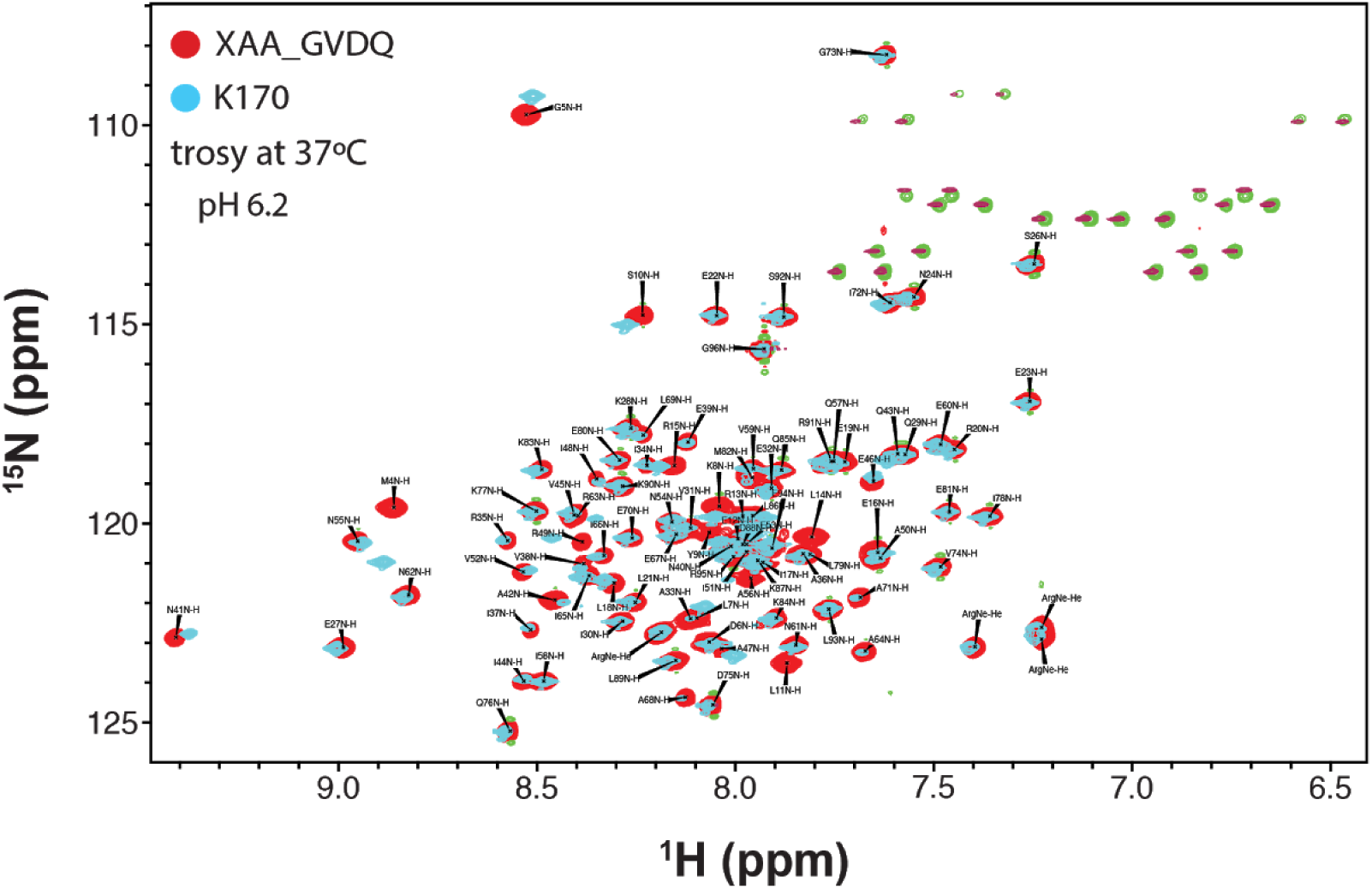
NMR assignments for XAA_GVDQ and XAA_GVDQ mutant M4L. TROSY showing that XAA_GVDQ and k170 (a single point mutant of XAA_GVDQ with M4L designed to facilitate NMR experiments) show the two structures are identical except at the M4 and G5.

### Supplemental Tables

**Table S1.**
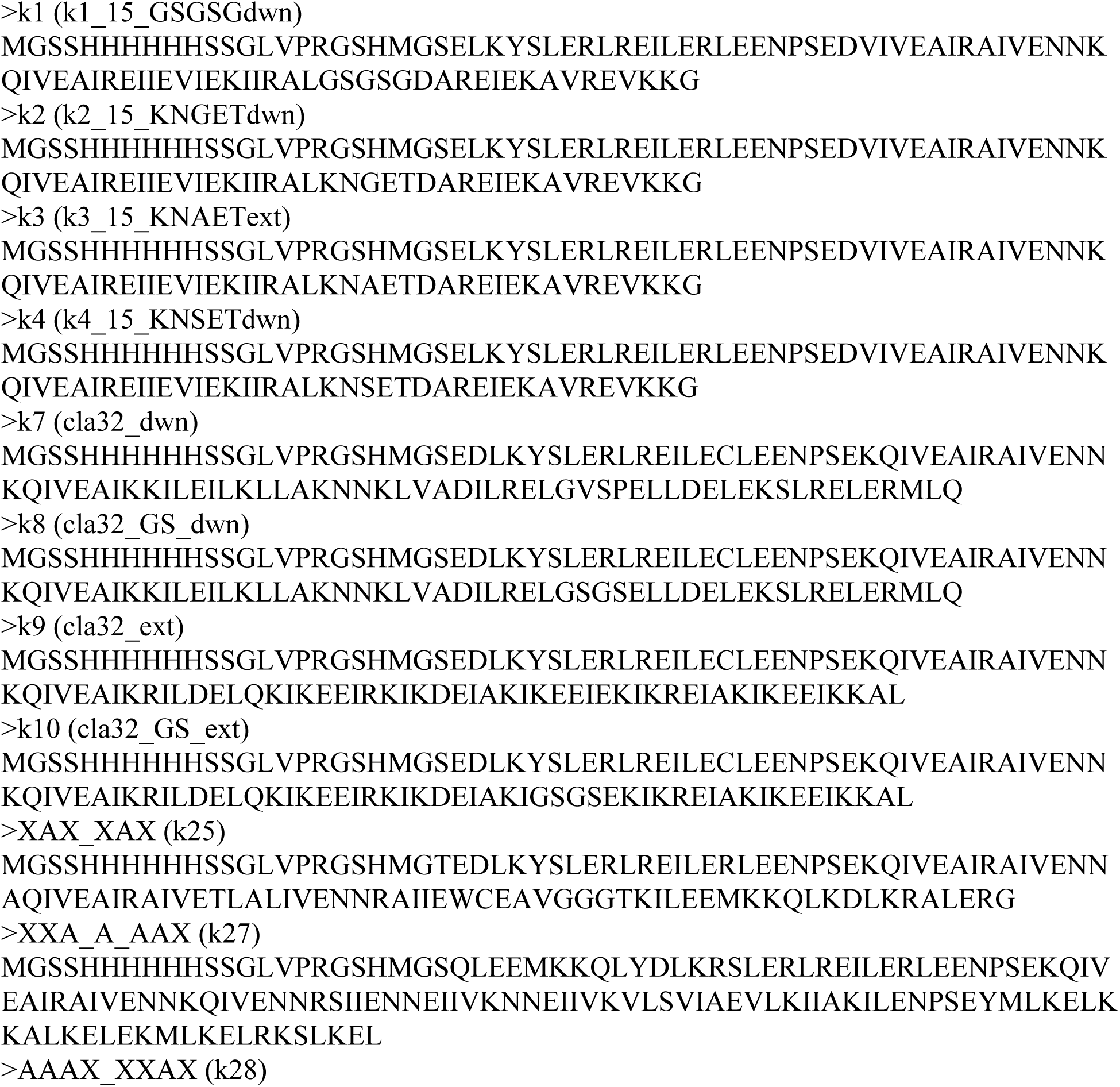
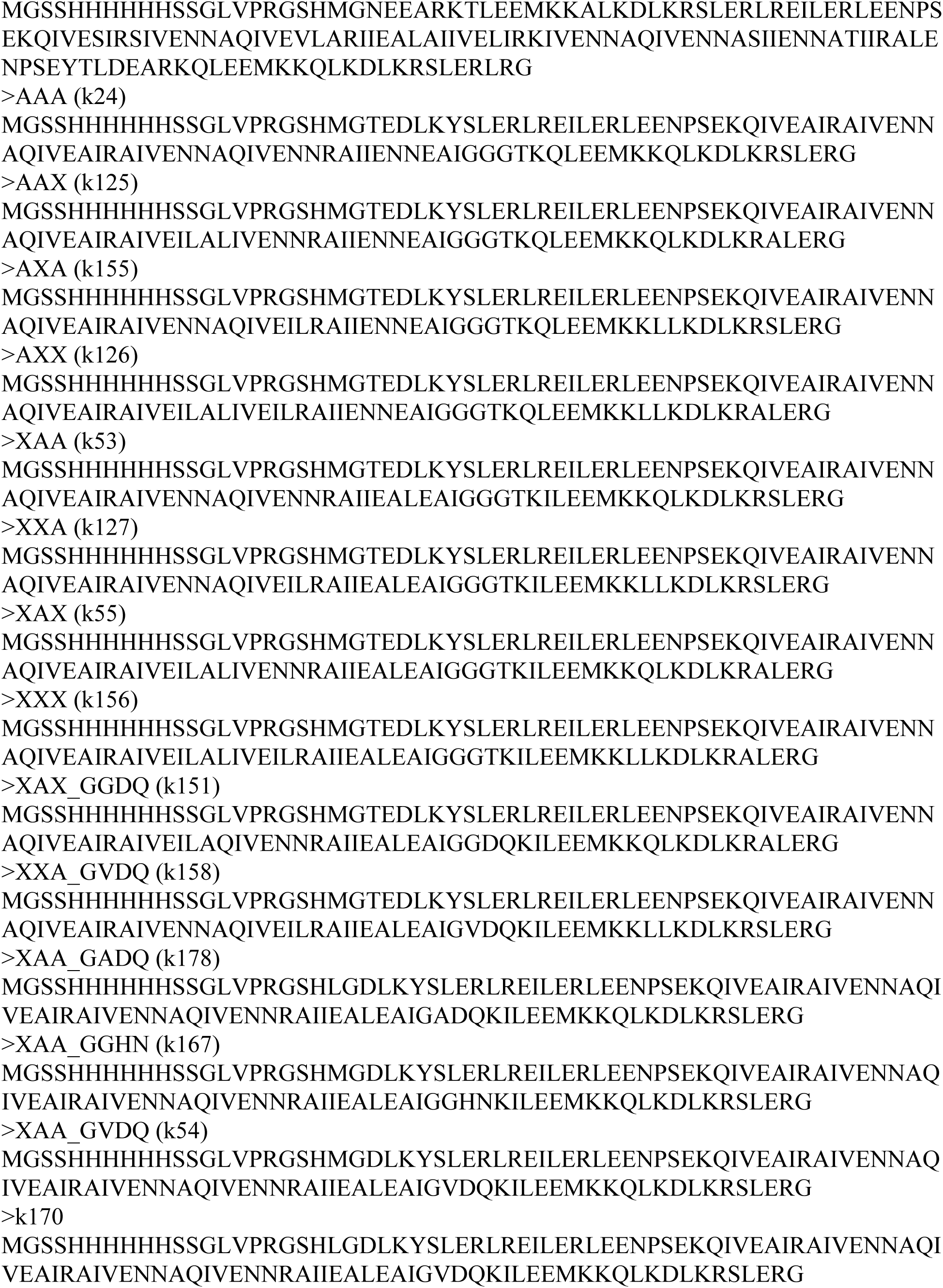
Amino acid sequence of protein designs tested in fasta format. Rosetta designs begin two residues after the thrombin cleavage and NdeI cut site of the pET28b backbone (LVPRGSHM). The glycine after the NdeI cut site is introduced manually to provide flexible linkage to the preceding histidine tag. Thrombin cleavage removes the N-terminal tag up the the apostrophe in the sequence LVPR’GS. k170 is a point mutant of XAA_GVDQ designed to remove the methionine from the plasmid backbone to facilitate NMR characterization and behaves identically to XAA_GVDQ in NMR experiments (Fig. S15). Alternative names are given in parenthesis)

**Table S2.** Protein crystallization conditions of structures reported.

| **Design Name** | **Screen** | **Crystallization condition** | **Temp** |
| --- | --- | --- | --- |
| AAA | Morpheus B10 | 10% w/v PEG 8000, 20% v/v ethylene glycol, 0.03 M of each halide (sodium fluoride, sodium bromide, sodium iodide), 0.1 M bicine/Trizma base pH 8.5 | 4°C |
| XXA | JCSG Core III E7 | 0.16 M magnesium acetate, 0.08 M sodium cacodylate pH 6.5, 20% (v/v) glycerol | 18°C |
| XAX | JCSG Core I C1 | 0.2 M ammonium acetate, 20% (w/v) PEG 3350 | 4°C |
| XAX_GGDQ | JCSG Core II E3 | 1.0 M lithium chloride, 0.1 M MES pH 6.0, 20% (w/v) PEG 6000 | 18°C |
| XXA_GVDQ | Morpheus A3 | 10% w/v PEG 4000, 20% v/v glycerol, 0.03 M of each divalent cation (magnesium chloride, calcium chloride), 0.1 M MES/imidazole pH 6.5 | 18°C |
| XAA_GGHN | JCSG Core IV A3 | 0.1 M CAPS pH 10.5, 40% (v/v) MPD | 18°C |
| XAA_GVDQ | JCSG Core III H1 | 2.0 M sodium formate, 0.1 M sodium acetate pH 4.6 | 18°C |
| XAA_GVDQ_calcium | JCSG Core IV G12 | 0.05 M calcium acetate, 0.1 M sodium acetate pH 4.5, 40% (w/v) 1,2-propanediol | 18°C |

**Table S3.** X-ray crystallography data collection and refinement statistics. Except for AAA (sent frozen to ALS 8.2.1), all other crystals were looped and frozen at the beamline (ALS 8.3.1). Beamline recommended strategies for collection was used. All structures suffered from poor diffraction data, some degree of radiation damage, potentially incorrect space group determination, and/or tNCS, resulting in higher R-free than typically expected of 1.9-2.8 Å resolution datasets. In regards to space group, automatic processing by XDS almost always determined the space group to be H32, which was incorrect. Reprocessing in other space groups (C2, P2, and P1) was tested, but not exhaustively.

| **Design Name**<br>**Biological assembly**<br>**PDB ID** | **AAA**<br>**trimer**<br>**6NX2** | **XXA**<br>**trimer**<br>**6NYI** | **XAX**<br>**trimer**<br>**6NYE** |
| --- | --- | --- | --- |
| Resolution range | 35.93 - 2.303 (2.385 - 2.303) | 63.18 - 2.3 (2.382 - 2.3) | 48.47 - 1.9 (1.968 - 1.9) |
| Space group | C 1 2 1 | C 1 2 1 | P 21 21 21 |
| Unit cell | 89.592 51.65 85.527 90 110.485 90 | 129.403 51.213 48.838 90 102.447 90 | 35.179 75.322 96.944 90 90 90 |
| Total reflections | 58747 (4463) | 93511 (8762) | 251950 (15068) |
| Unique reflections | 16161 (1486) | 14041 (1384) | 21012 (2046) |
| Multiplicity | 3.6 (3.0) | 6.7 (6.3) | 12.0 (7.4) |
| Completeness (%) | 90.14 (69.63) | 92.10 (82.28) | 96.77 (87.78) |
| Mean I/sigma(I) | 9.42 (1.32) | 10.01 (1.29) | 14.25 (0.85) |
| Wilson B-factor | 55.42 | 32.95 | 26.59 |
| R-merge | 0.0686 (0.7771) | 0.137 (1.532) | 0.1199 (1.974) |
| R-meas | 0.08064 (0.9321) | 0.1487 (1.67) | 0.1252 (2.125) |
| R-pim | 0.04202 (0.5077) | 0.05723 (0.6565) | 0.03571 (0.768) |
| CC1/2 | 0.996 (0.709) | 0.999 (0.737) | 0.999 (0.378) |
| CC* | 0.999 (0.911) | 1 (0.921) | 1 (0.74) |
| Reflections used in refinement | 16161 (1137) | 14041 (1142) | 21012 (1818) |
| Reflections used for R-free | 1487 (111) | 1312 (114) | 1703 (150) |
| R-work | 0.2594 (0.3944) | 0.2588 (0.4285) | 0.2019 (0.3513) |
| R-free | 0.2791 (0.4538) | 0.2810 (0.4488) | 0.2335 (0.3626) |
| CC(work) | 0.942 (0.746) | 0.967 (0.817) | 0.971 (0.668) |
| CC(free) | 0.927 (0.663) | 0.958 (0.602) | 0.932 (0.637) |
| Number of non-hydrogen atoms | 1957 | 2107 | 2317 |
| macromolecules | 1944 | 2077 | 2141 |
| ligands | 2 |  |  |
| solvent | 11 | 30 | 176 |
| Protein residues | 267 | 286 | 283 |
| RMS(bonds) | 0.005 | 0.004 | 0.005 |
| RMS(angles) | 0.51 | 0.79 | 0.81 |
| Ramachandran favored (%) | 99.62 | 98.19 | 100 |
| Ramachandran allowed (%) | 0.38 | 1.81 | 0 |
| Ramachandran outliers (%) | 0 | 0 | 0 |
| Rotamer outliers (%) | 6.86 | 1.04 | 0.97 |
| Clashscore | 1.61 | 4.33 | 2.08 |
| Average B-factor | 88.09 | 51.87 | 31.79 |
| macromolecules | 88.14 | 51.91 | 31.42 |
| ligands | 68.44 |  |  |
| solvent | 82.61 | 49.08 | 36.26 |
| Number of TLS groups | 3 |  |  |

| Design Name | XAX_GGDQ | XXA_GVDQ | XAA_GGHN |
| --- | --- | --- | --- |
| Biological assembly | trimer | trimer | trimer |
| PDB ID | 6NYK | 6NZ1 | 6NZ3 |
| Resolution range | 45.34 - 2.8 (2.9 - 2.8) | 88.36 - 1.9 (1.968 - 1.9) | 43.67 - 2.3 (2.382 - 2.3) |
| Space group | C 1 2 1 | P 1 21 1 | C 1 2 1 |
| Unit cell | 95.416 55.002 57.906 90 123.381 90 | 50.647 76.747 91.946 90 106.064 90 | 88.741 51.271 85.873 90 110.076 90 |
| Total reflections | 41579 (4425) | 355141 (36439) | 107072 (10366) |
| Unique reflections | 6209 (631) | 53261 (5304) | 16278 (1626) |
| Multiplicity | 6.7 (7.0) | 6.7 (6.9) | 6.6 (6.4) |
| Completeness (%) | 95.12 (90.73) | 99.70 (99.77) | 98.15 (96.20) |
| Mean I/sigma(I) | 10.20 (1.40) | 8.98 (1.46) | 31.70 (8.63) |
| Wilson B-factor | 82.44 |  | 47.9 |
| R-merge | 0.08044 (1.306) | 0.1542 (1.073) | 0.03 (0.1729) |
| R-meas | 0.08715 (1.409) | 0.168 (1.161) | 0.03272 (0.1885) |
| R-pim | 0.03319 (0.5248) | 0.06572 (0.4397) | 0.01288 (0.07428) |
| CC1/2 | 0.999 (0.609) | 0.995 (0.703) | 1 (0.996) |
| CC* | 1 (0.87) | 0.999 (0.908) | 1 (0.999) |
| Reflections used in refinement | 6209 (587) | 53261 (5304) | 16278 (1571) |
| Reflections used for R-free | 607 (46) | 1563 (162) | 926 (93) |
| R-work | 0.2617 (0.2997) | 0.1847 (0.3347) | 0.2620 (0.2687) |
| R-free | 0.2971 (0.2876) | 0.2159 (0.3631) | 0.3017 (0.3071) |
| CC(work) | 0.974 (0.760) | 0.883 (0.697) | 0.961 (0.911) |
| CC(free) | 0.944 (0.877) | 0.839 (0.413) | 0.917 (0.793) |
| Number of non-hydrogen atoms | 1729 | 4574 | 2041 |
| macromolecules | 1729 | 4317 | 2030 |
| ligands |  |  | 1 |
| solvent |  | 257 | 10 |
| Protein residues | 271 | 569 | 275 |
| RMS(bonds) | 0.002 | 0.009 | 0.004 |
| RMS(angles) | 0.41 | 0.9 | 0.54 |
| Ramachandran favored (%) | 98.85 | 97.49 | 99.63 |
| Ramachandran allowed (%) | 1.15 | 2.51 | 0.37 |
| Ramachandran outliers (%) | 0 | 0 | 0 |
| Rotamer outliers (%) | 0.86 | 0.71 | 0 |
| Clashscore | 6.65 | 6.1 | 6.27 |
| Average B-factor | 79.97 | 29.48 | 63.99 |
| macromolecules | 79.97 | 29.27 | 64.01 |
| ligands |  |  | 56.69 |
| solvent |  | 33.13 | 60.62 |
| Number of TLS groups |  | 17 |  |

| **Design Name** | **XAA_GVDQ** | **XAA_GVDQ_calcium** |
| --- | --- | --- |
| **Biological assembly** | **hexamer** | **trimer** |
| **PDB ID** | **6NXM** | **6NY8** |
| Resolution range | 47.71 - 2.2 (2.279 - 2.2) | 80.05 - 2.3 (2.382 - 2.3) |
| Space group | C 1 2 1 | C 1 2 1 |
| Unit cell | 101.045 58.282 58.793 90 125.046 90 | 86.158 49.602 85.032 90 109.713 90 |
| Total reflections | 95755 (9491) | 98944 (9361) |
| Unique reflections | 14333 (1411) | 15110 (1505) |
| Multiplicity | 6.7 (6.7) | 6.5 (6.2) |
| Completeness (%) | 96.43 (89.65) | 94.59 (85.71) |
| Mean I/sigma(I) | 25.02 (2.53) | 21.93 (1.90) |
| Wilson B-factor | 50.43 | 55.74 |
| R-merge | 0.03291 (0.6889) | 0.03765 (1.059) |
| R-meas | 0.0358 (0.7469) | 0.041 (1.156) |
| R-pim | 0.01389 (0.2855) | 0.01598 (0.457) |
| CC1/2 | 1 (0.905) | 1 (0.864) |
| CC* | 1 (0.975) | 1 (0.963) |
| Reflections used in refinement | 14333 (1265) | 15110 (1308) |
| Reflections used for R-free | 1202 (109) | 1235 (111) |
| R-work | 0.2457 (0.3578) | 0.2875 (0.4163) |
| R-free | 0.2770 (0.4159) | 0.3041 (0.4535) |
| CC(work) | 0.977 (0.806) | 0.948 (0.858) |
| CC(free) | 0.940 (0.681) | 0.905 (0.842) |
| Number of non-hydrogen atoms | 1874 | 1917 |
| macromolecules | 1868 | 1915 |
| ligands |  | 2 |
| solvent | 6 |  |
| Protein residues | 274 | 272 |
| RMS(bonds) | 0.004 | 0.004 |
| RMS(angles) | 0.83 | 0.77 |
| Ramachandran favored (%) | 98.51 | 98.12 |
| Ramachandran allowed (%) | 1.49 | 1.88 |
| Ramachandran outliers (%) | 0 | 0 |
| Rotamer outliers (%) | 0.69 | 1.23 |
| Clashscore | 3.34 | 0 |
| Average B-factor | 72.91 | 79.41 |
| macromolecules | 72.93 | 79.41 |
| ligands |  | 80.45 |
| solvent | 68.55 |  |

**Table S4.** SAXS molecular weight measurements for select designs.

| **Design** | **Expected trimer (MW)** | **SAXS condition** | **MW** |
| --- | --- | --- | --- |
| XAX | 33138 | pH 7 | 34000 |
|  |  | pH 5 | 39000 |
| XAA_GVDQ | 32938 | 0 mM calcium | 28000 |
|  |  | 15 mM calcium | 29000 |
|  |  | 20 mM calcium | 31000 |
|  |  | 30 mM calcium | 32000 |

**Table S5.**
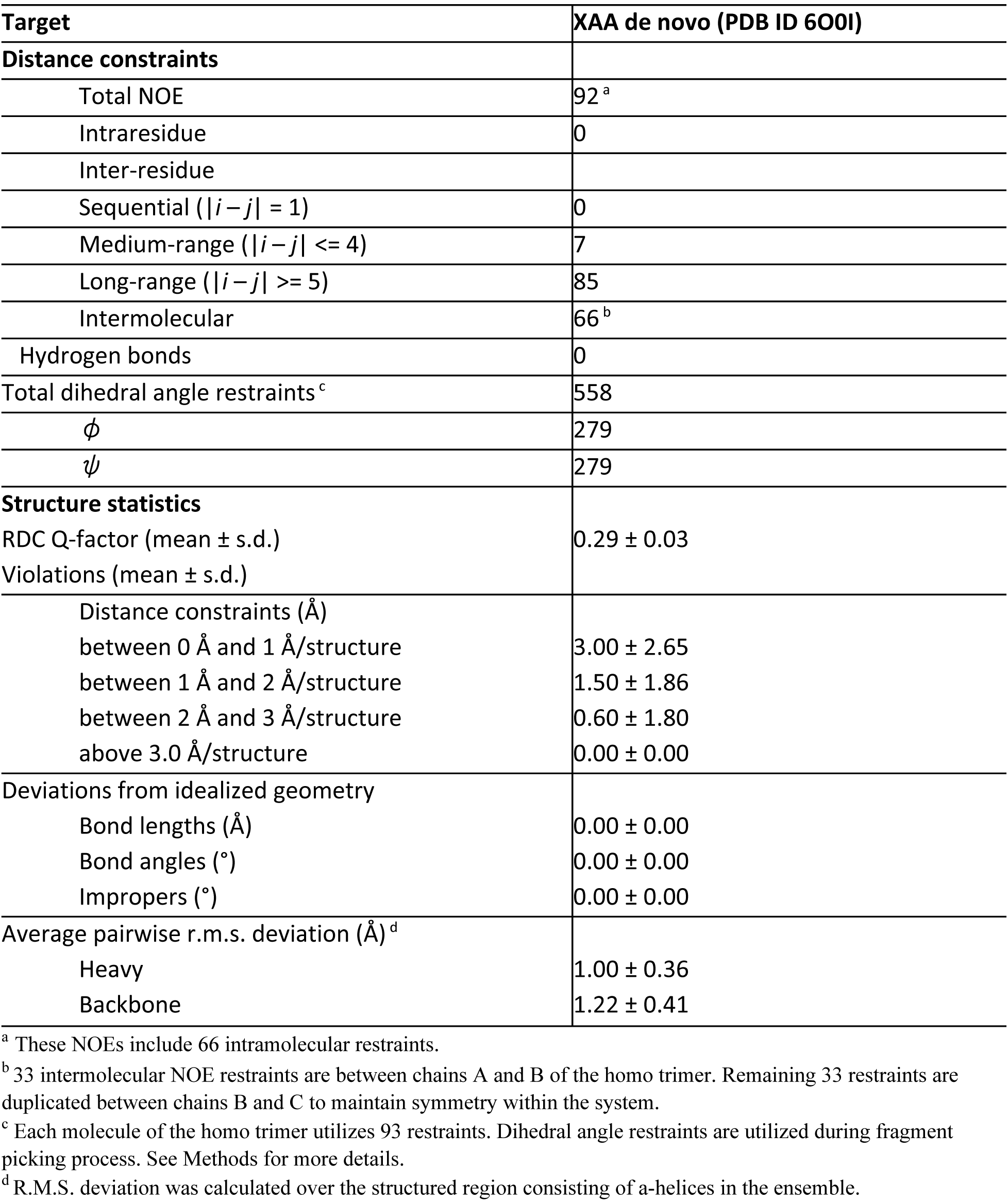
NMR restraints and structural statistics for XAA structural ensembles. A set of structure calculation for XAA *de novo* using Chemical Shifts, NOEs and RDC data was performed.

| Target | XAA de novo (PDB ID 6O0I) |
| --- | --- |
| **Distance constraints** |  |
| Total NOE | 92 <sup>a</sup> |
| Intraresidue | 0 |
| Inter-residue |  |
| Sequential ($\vert i - j\vert = 1$) | 0 |
| Medium-range ($\vert i - j\vert \leq 4$) | 7 |
| Long-range ($\vert i - j\vert \geq 5$) | 85 |
| Intermolecular | 66 <sup>b</sup> |
| Hydrogen bonds | 0 |
| Total dihedral angle restraints <sup>c</sup> | 558 |
| $\phi$ | 279 |
| $\psi$ | 279 |
| **Structure statistics** |  |
| RDC Q-factor (mean ± s.d.) | 0.29 ± 0.03 |
| Violations (mean ± s.d.) | |
| Distance constraints (Å) |  |
| between 0 Å and 1 Å/structure | 3.00 ± 2.65 |
| between 1 Å and 2 Å/structure | 1.50 ± 1.86 |
| between 2 Å and 3 Å/structure | 0.60 ± 1.80 |
| above 3.0 Å/structure | 0.00 ± 0.00 |
| Deviations from idealized geometry |  |
| Bond lengths (Å) | 0.00 ± 0.00 |
| Bond angles (°) | 0.00 ± 0.00 |
| Impropers (°) | 0.00 ± 0.00 |
| Average pairwise r.m.s. deviation (Å) <sup>d</sup> |  |
| Heavy | 1.00 ± 0.36 |
| Backbone | 1.22 ± 0.41 |

<sup>a</sup> These NOEs include 66 intramolecular restraints.
<sup>b</sup> 33 intermolecular NOE restraints are between chains A and B of the homo trimer. Remaining 33 restraints are duplicated between chains B and C to maintain symmetry within the system.
<sup>c</sup> Each molecule of the homo trimer utilizes 93 restraints. Dihedral angle restraints are utilized during fragment picking process. See Methods for more details.
<sup>d</sup> R.M.S. deviation was calculated over the structured region consisting of $\alpha$ -helices in the ensemble.

**Table S6.** NMR restraints and structural statistics for k170 structural ensembles. Three sets of structure calculations for k170 (i) without calcium *de novo* using Chemical Shifts, NOEs and RDC data, (ii) without calcium using NMR Chemical Shifts and RDC data, and (iii) with calcium using NMR Chemical Shifts and RDC data were performed. k170 is a point mutant of XAA_GVDQ that removes a methionine from the plasmid backbone and behaves identically to XAA_GVDQ in NMR experiments (Fig. S15).

| Target | k170 *de novo* without calcium (PDB ID 6O0C) | k170 without calcium | k170 with calcium |
| --- | --- | --- | --- |
| **Distance constraints** |  |  |  |
| Total NOE | 139<sup>a</sup> | 0 | 0 |
| Intraresidue | 0 | 0 | 0 |
| Inter-residue |  |  |  |
| Sequential ($\vert i - j\vert = 1$) | 0 | 0 | 0 |
| Medium-range ($\vert i - j\vert \leq 4$) | 23 | 0 | 0 |
| Long-range ($\vert i - j\vert \geq 5$) | 116 | 0 | 0 |
| Intermolecular | 56<sup>b</sup> | 0 | 0 |
| Hydrogen bonds | 0 | 0 | 0 |
| Total dihedral angle restraints<sup>c</sup> | 528 | 528 | 528 |
| $\phi$ | 264 | 264 | 264 |
| $\psi$ | 264 | 264 | 264 |
| **Structure statistics** |  |  |  |
| RDC Q-factor (mean ± s.d.) | 0.18 ± 0.01 | 0.21 ± 0.01 | 0.21 ± 0.01 |
| Violations (mean ± s.d.) | | | |
| Distance constraints (Å) |  |  |  |
| between 0 Å and 1 Å/structure | 1.80 ± 1.80 | 0.00 ± 0.00 | 0.00 ± 0.00 |
| between 1 Å and 2 Å/structure | 0.30 ± 0.50 | 0.00 ± 0.00 | 0.00 ± 0.00 |
| above 2.0 Å/structure | 0.00 ± 0.00 | 0.00 ± 0.00 | 0.00 ± 0.00 |
| Deviations from idealized geometry |  |  |  |
| Bond lengths (Å) | 0.00 ± 0.00 | 0.00 ± 0.00 | 0.00 ± 0.00 |
| Bond angles (°) | 0.00 ± 0.00 | 0.00 ± 0.00 | 0.00 ± 0.00 |
| Impropers (°) | 0.00 ± 0.00 | 0.00 ± 0.00 | 0.00 ± 0.00 |
| Average pairwise r.m.s. deviation (Å)<sup>d</sup> |  |  |  |
| Heavy | 0.88 ± 0.20 | 0.52 ± 0.20 | 0.54 ± 0.21 |
| Backbone | 0.78 ± 0.17 | 0.47 ± 0.20 | 0.49 ± 0.23 |

<sup>a</sup> These NOEs include 83 intramolecular restraints.
<sup>b</sup> 28 intermolecular NOE restraints are between chains A and B of the homo trimer. Remaining 28 restraints are duplicated between chains B and C to maintain symmetry within the system.
<sup>c</sup> Each molecule of the homo trimer utilizes 88 restraints. Dihedral angle restraints are utilized during fragment picking process. See Methods for more details.
<sup>d</sup> R.M.S. deviation was calculated over the structured region consisting of $\alpha$ -helices in the ensemble.

### Supplemental Text

#### Tuning of the free energy difference between short and long states

Crystal structures of designs XXA (PDB ID 6nyi) and XAX (PDB ID 6nye) match the modeled short state with Cα root-mean-square deviations (RMSDs) of 1.37 and 0.97, respectively at 1.9-2.3 Å resolution (Fig. 2a,b). For both, most of the deviation, as expected, is in the flexible hinge: the crystal structures show that each monomer adopts a different loop. For XXA, the crystal structure confirms the short state, but there are deviations from the model at the hydrogen bond network, which is not surprisingly since it is in a solvent exposed position. For XAX, the designed hydrogen bond network formed as expected (Fig 2b. “network A” panel vs. Fig. 2a “network A” panel). To further characterize the solution structure, we prepared a selective AI(LV)^proS^ methyl-labeled XAX sample. The 3D C_M_-C_M_H_M_ NOESY spectrum for XAX shows NOE patterns consistent with the compact structure, with many unambiguously assigned close-range NOEs identified between the ^13^Cδ1 methyl of Ile66 and ^13^Cδ2 of Leu86 (Fig. 2c).

Attempts to solve a crystal structure for XAA were unsuccessful, but a ^15^N MAI(LV)^proS^ methyl-labeled sample prepared for the protein showed highly dispersed NMR spectra. The fully assigned methyl HMQC spectra show resonances of increased linewidth, relative to our previously assigned XAX construct, indicating the presence of dynamics at the μsec-msec timescale. The fully assigned backbone amide TROSY-HSQC spectra confirm this result, and further reveal the presence of a minor component (∼10-15% population, not exchanging with the major component at the NMR timescale, as confirmed by TROSY ZZ exchange experiments) for residues distributed at the interface between the inner and outer helices, (Fig. S14), likely due to local breaking of the regular C3 symmetry. Comparison of the shared methyl NOEs between the major state of XAA and XAA_GVDQ (which by NMR is locked in the short state) (Fig. S8a) and analysis of Residual Dipolar Couplings (RDCs) (Fig. S8b) further suggest that the flipping helix is in the short state, adopting an alternative packing against the inner helices. Finally, a comparison with the *de novo* NMR structure (solved using chemical shifts, long-range NOEs and RDCs; Table S5) clearly shows that the flipping helix is shifted by about a turn up towards the hinge (Fig. 2d,e). The C-terminal hydrogen bond network in XAA is solvent exposed and more likely to be disrupted by competing water interactions. This disrupted network likely destabilizes the interface between the flipping helix and the inner helices, allowing alternative packing. In contrast, the C-terminal region of XAX is hydrophobic, very well packed in the crystal structure and effectively stabilizes the flipping helix.

Design AAA (PDB ID 6nx2) adopts the long state according to crystallographic evidence with RMSD = 1.40 at about 2.3 Å resolution (Fig. 2e). While the helical propensity of the residues at and surrounding the hinge region was predicted to be low, in the crystal structure this region is clearly helical and the most flexible residues are at the C-terminus (Fig. S14). The structure of AAA deviates from the predicted model because the asparagines of two of the hydrogen bond networks coordinate bromide ions (ions and waters are not explicitly accounted for during modeling). In particular, N61 shifts toward the center of the helix (relative to the model to coordinate the ion (Fig. S10). Taken together, there is potential for a single sequence that can interconvert between the two designed states since a change in three residues, corresponding to the location of a hydrogen bond network, can switch the observed state (i.e., XAA vs AAA).

